# siRNA-mediated gene knockdown via electroporation in hydrozoan jellyfish embryos

**DOI:** 10.1101/2022.03.24.485716

**Authors:** Tokiha Masuda-Ozawa, Sosuke Fujita, Ryotaro Nakamura, Hiroshi Watanabe, Erina Kuranaga, Yu-ichiro Nakajima

## Abstract

As the sister group to bilaterians, cnidarians stand in a unique phylogenetic position that provides insight into evolutionary aspects of animal development, physiology, and behavior. While cnidarians are classified into two types, sessile polyps and free-swimming medusae, most studies at the cellular and molecular levels have been conducted on representative polyp-type cnidarians and have focused on establishing techniques of genetic manipulation. Recently, gene knockdown by delivery of short hairpin RNAs into eggs via electroporation has been introduced in two polyp-type cnidarians, *Nematostella vectensis* and *Hydractinia symbiolongicarpus*, enabling systematic loss-of-function experiments. By contrast, current methods of genetic manipulation for most medusa-type cnidarians, or jellyfish, are quite limited, except for *Clytia hemisphaerica*, and reliable techniques are required to interrogate function of specific genes in different jellyfish species. Here, we present a method to knock down target genes by delivering small interfering RNA (siRNA) into fertilized eggs via electroporation, using the hydrozoan jellyfish, *Clytia hemisphaerica* and *Cladonema paciificum*. We show that siRNAs targeting endogenous *GFP1* and *Wnt3* in *Clytia* efficiently knock down gene expression and result in known planula phenotypes: loss of green fluorescence and defects in axial patterning, respectively. We also successfully knock down endogenous *Wnt3* in *Cladonema* by siRNA electroporation, which circumvents the technical difficulty of microinjecting small eggs. *Wnt3* knockdown in *Cladonema* causes gene expression changes in axial markers, suggesting a conserved Wnt/β-catenin-mediated pathway that controls axial polarity during embryogenesis. Our gene-targeting siRNA electroporation method is applicable to other animals, including and beyond jellyfish species, and will facilitate the investigation and understanding of myriad aspects of animal development.

## Introduction

The phylum Cnidaria is the sister group to Bilateria, having separated from their common ancestor over 500 million years ago. Cnidarians have diversified their morphologies across an array forms, including corals, sea anemones, hydroids, and jellyfish, all of which are divided into two clades: Anthozoa and Medusozoa. The two differ in that Anthozoa includes only sessile polyp-type animals, while Medusozoa (Hydrozoa, Staurozoa, Scyphozoa, and Cubozoa) contains two forms: polyp and medusa, commonly known as jellyfish (Fig. 1A)^1,2^. The life cycle of most, though not all, jellyfish consists of five forms: gametes, fertilized eggs, planulae, polyps, and medusae^3^. Fertilized eggs undergo embryogenesis to become planula larvae, which metamorphose into sessile polyps. While vegetatively-growing polyps give rise to free-swimming medusae through the process of budding or strobilization, medusae sexually reproduce by releasing gametes. Due to their unique phylogenetic position, studies using cnidarians have provided evolutionary insight into development, regeneration, and behaviors in multicellular animals^1,4,5^. Despite divergent morphologies and lifestyles among cnidarians, to date, the molecular and cellular understanding of cnidarians has been acquired primarily using polyp-type animals such as the anthozoan *Nematostella vectensis* and the hydrozoan *Hydra* and *Hydractinia*. By contrast, jellyfish biology remains largely unestablished.

**Figure 1.**
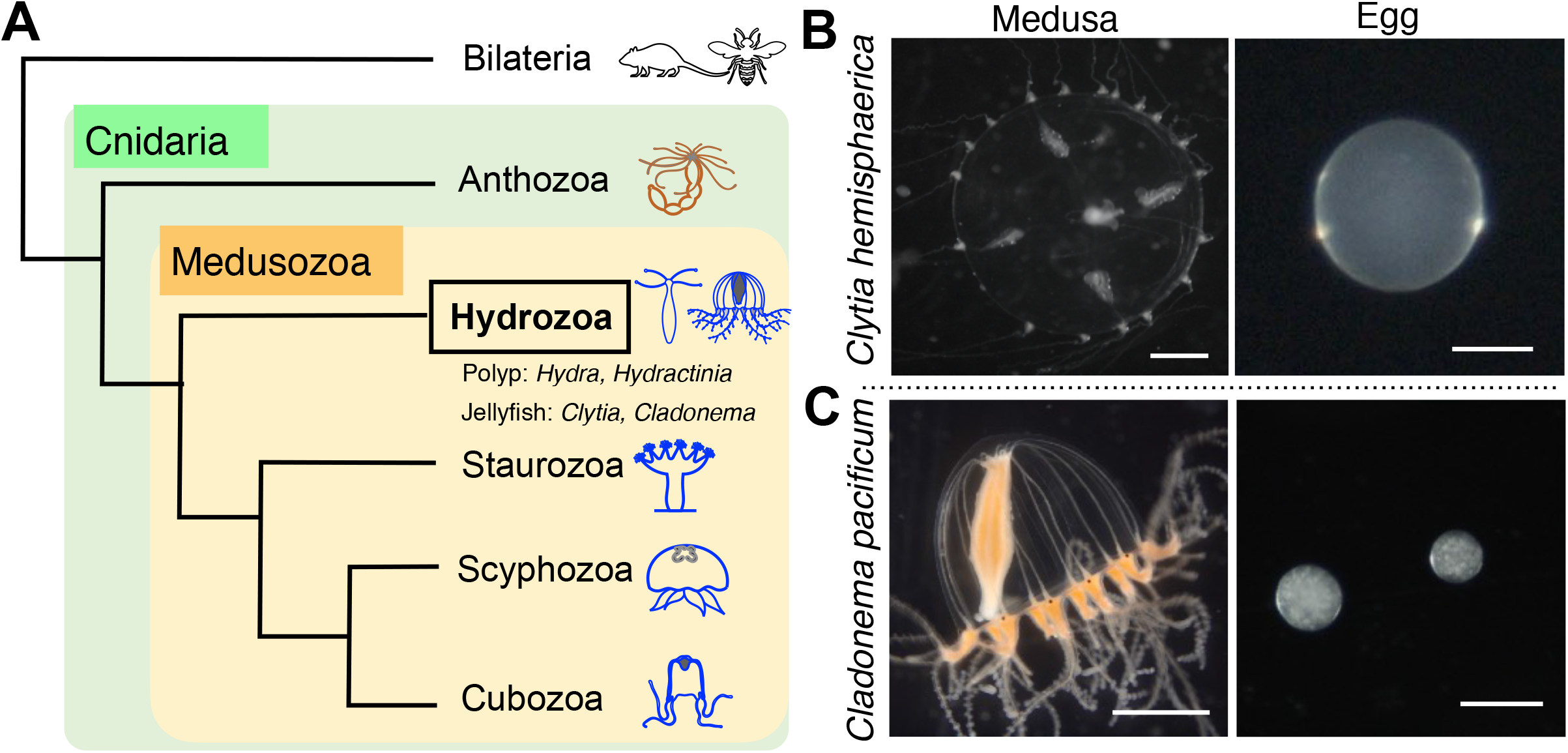
The hydrozoan jellyfish *Clytia hemisphaerica* and *Cladonema pacificum*. (**A**) Cladogram depicting the phylogenetic position of cnidarian jellyfish. As the sister group of Bilateria, the phylum Cnidaria is divided into two clades, Anthozoa and Medusozoa, which consists of four classes: Hydrozoa, Staurozoa, Scyphozoa, and Cubozoa. Hydrozoa includes polyp type animals without a medusa stage (e.g. *Hydra* and *Hydractinia)* and jellyfish that have both polyp and medusa stages (e.g. *Clytia* and *Cladonema*). (**B** and **C**) Photos of an adult medusa and the eggs of *Clytia hemisphaerica* (**B**) and *Cladonema pacificum* (**C**). Scale bars: 1 mm for medusae; 100 μm for eggs.

Among jellyfish species, the hydrozoan *Clytia hemisphaerica* is the best-studied laboratory model^6,7^. The well-established methodology for maintaining and propagating animals ensures a reliable source for daily experiments^8^. Transparent and small-sized medusae make whole-mount visualization achievable, and relatively large-sized eggs (~180 μm diameter) enable different manipulations via microinjection (Fig. 1B). Indeed, gene silencing by morpholino oligos and mRNA microinjection as well as gene knockout via CRISPR/Cas9 allow for the investigation of functions of genes of interest^9–13^. Furthermore, the *Clytia* genome assembly is complete, and transcriptome profiles are available for various stages and tissues, even at the single-cell level, which creates the foundation for research at the molecular level^12,14–18^. These resources and techniques, together with recently-established transgenesis^19^, have accelerated research into numerous facets of distinct *Clytia* life stages. While initial studies focused on embryogenesis, current research topics using *Clytia* include gametogenesis, regeneration, and behavior exhibited by adult medusa ^9,10,12,17,19,20^.

Compared to the established model jellyfish *Clytia*, the hydrozoan *Cladonema pacificum* is an emerging model that has recently been utilized for various studies in development, regeneration, and physiology^21^. The ease of rearing all stages of *Cladonema* without a filtration system or large water tank, along with their high spawning rate, enable easy lab maintenance. Small-sized medusae with branched tentacles allow for investigations of body size control and tentacle morphogenesis (Fig. 1C)^22–24^. *Cladonema* gametogenesis is regulated by a light-dark cycle as in *Clytia*, and recent work has identified the neuropeptides involved in oocyte maturation^25^. Furthermore, *Cladonema* medusae possess eyes with a complex structure that includes lenses (ocelli) in their tentacle bulbs, and studies using the closely related species *Cladonema radiatum* have identified conserved light-sensitive opsins and regulators of eye formation (*Pax*, *Six*, and *Eya*)^25–29^, providing a model for the evolutionary developmental biology of photoreceptor organs. Despite these attractive features of *Cladonema* as a laboratory animal, no genome assembly or transcriptome is currently available, and genetic manipulation techniques are only just being developed. One major technical issue in manipulating *Cladonema* is their small eggs—with a diameter of approximately 60 μm (Fig. 1C), regular microinjection is quite difficult. Establishing genetic manipulations is required to facilitate the in-depth investigations needed to further understand the biology of *Cladonema*, and an alternative method that eliminates the need for microinjection would greatly facilitate that objective.

RNA interference (RNAi), the phenomenon of double stranded RNA (dsRNA) mediated silencing of target genes, has been widely exploited in living organisms to analyze gene function^30^, primarily using chemically- or *in vitro*-synthesized doublestranded small interfering RNAs (siRNAs) or vector-based short hairpin RNAs (shRNAs)^31^. In cnidarians, siRNA-mediated gene silencing was initially applied to *Nematostella* and *Hydra* polyps via soaking or electroporation^32–34^. More recently, gene knockdown via shRNA microinjection or electroporation into eggs has been utilized in studies with *Nematostella* and *Hydractinia* to show efficient reduction in gene expression and associated phenotypes in early developmental stages^35–38^. In particular, shRNA delivery via electroporation, which does not require the rigors of microinjection, allows for the experimental gene knockdown of large numbers of individuals simultaneously^36,38^, opening up the possibility of manipulating genes in different aquatic animals, even those that produce very small eggs like *Cladonema*.

The mechanism of RNAi-mediated gene repression after introducing foreign dsRNA into cells involves the microRNA (miRNA) pathway, an endogenous gene repression machinery conserved in both animals and plants^39^. In cnidarians, the presence of the miRNA pathway has been demonstrated in *Nematostella* and *Hydra*^32,40,41^. However, little is known about the endogenous RNAi pathway, especially the presence and roles of miRNAs and the miRNA-related genes, in jellyfish such as *Clytia* and *Cladonema*. Furthermore, in mammalian cultured cells, siRNAs can induce gene repression independent of endogenous dsRNA processing factors, while shRNAs cannot^42,43^. On this basis, we selected siRNA instead of shRNA for gene knockdown to avoid the potential pitfall of dsRNA processing in jellyfish.

Here we report a gene knockdown method for jellyfish embryos with siRNA via electroporation. Using two hydrozoan species, *Clytia hemisphaerica* and *Cladonema pacificum*, we demonstrate that siRNA delivery into fertilized eggs effectively reduces the expression of endogenous genes. We also confirm the known loss-of-function phenotypes in *Clytia* after knocking down *GFP1* or *Wnt3*, which are induced in a dose-dependent manner with siRNA. We further show that knockdown of *Wnt3* in *Cladonema* embryos results in the reduction of gene expression of the oral marker *Brachyury* in planula, implicating the Wnt/β-catenin pathway in the control of oral-aboral patterning. Overall, our siRNA-mediated knockdown approach allows for the manipulation of a large number of embryos through electroporation and enables functional analysis of early development, providing a new experimental platform applicable to different jellyfish species and other marine invertebrates.

## Materials and Methods

### Animal culture and spawning induction

*Clytia hemisphaerica* (Z11, Z4B as females and Z4C2, Z23 as males) and *Cladonema pacificum* (6W as females and UN2 as males) were used for this research.

*Clytia hemisphaerica* were cultured using a previously reported method^8^ with a few modifications. Artificial sea water (ASW) was prepared using 220 g SEA LIFE (Marin Tech) per 5L MilliQ water (Merk Millipore) with antibiotics (40 units/ml of penicillin and 40 μg/ml of streptomycin). Medusae, embryos, and planula larvae were maintained at 20°C. Medusae were fed daily with Vietnamese brine shrimp (A&A Marine LLC, Elk Rapids, MI, USA). Spawning timing was controlled by a 13 hours (h) dark /11 h light cycle, and spawning was induced by light. Male and female medusae were transferred into V-7 cups (AS ONE) before spawning (60 min for male and 90 min for female after light stimulation).

*Cladonema pacificum* were cultured as previously described^23^. *Cladonema* medusae were maintained at 22°C in ASW, which was prepared using SEA LIFE (Marin Tech) dissolved in tap water with chlorine neutralizer (Coroline off, GEX Co. ltd) (24 p.p.t) and antibiotics (40 units/ml penicillin and 40 μg/ml of streptomycin). Spawning timing was controlled by a 30 min dark/23.5 h light cycle, and spawning of male and female gametes was induced by dark stimulation. Before dark stimulation, adult females and males were separately transferred into V7 cups (AS ONE) and 60 mm dishes (BD), respectively.

*Nematostella vectensis* were cultured as previously described^44^, with a few modifications. Briefly, adult animals were maintained in brackish water at a salinity one-third of artificial seawater (35 g/l, pH 7.5-8.0, SEA LIFE, Marin Tech) and fed with freshly hatched artemia twice per week. Spawning induction was performed at 26°C under light for at least 11 h.

### siRNA and shRNA

siRNA sequences (19 mer RNA + 2 mer DNA) for *CheGFP1, CheWnt3*, and *CpWnt3* were designed based on their CDS sequences by Nippon Gene Co., Ltd, and siRNA duplexes were synthesized by manufacturers (Nippon Gene Co., Ltd and Sigma-Aldrich, Merk). Lyophilized siRNA was resuspended in RNase free water to a final concentration of 6 μg/μl as stock solution. The siRNA stock solution was diluted with RNase free water to a total volume of 10 μl and then added to 90 μl of fertilized eggs in 15% Ficoll ASW just prior to electroporation.

shRNAs were synthesized as described in previous reports^36,38^. Briefly, shRNAs were synthesized by *in vitro* transcription (IVT) from double stranded DNA templates using the AmpliScribeTM T7-flashTM transcription kit (Lucigen, Inc.) and were purified using Direct-Zol TM RNA Miniprep kits (Zymo Research, R2070). Concentrations of shRNA was measured with a NanoDrop One (Thermo fisher).

### Collection of gametes and electroporation for *Clytia* and *Cladonema* fertilized eggs

For the preparation of gametes, sexually-mature *Clytia* medusae were treated following a previously reported method^8^, partially modified to fit our experimental equipment. Adult male and female medusae were maintained on a 13 h dark/11 h light cycle, and light stimuli induced gametogenesis in 60 min for sperm and 90 min for eggs (Fig. 2A). For *Cladonema pacificum*, sexually-mature medusae were maintained on a 30 min dark/23.5 h light cycle. Dark stimulation induced gametogenesis in 25 min for both sexes.

**Figure 2.**
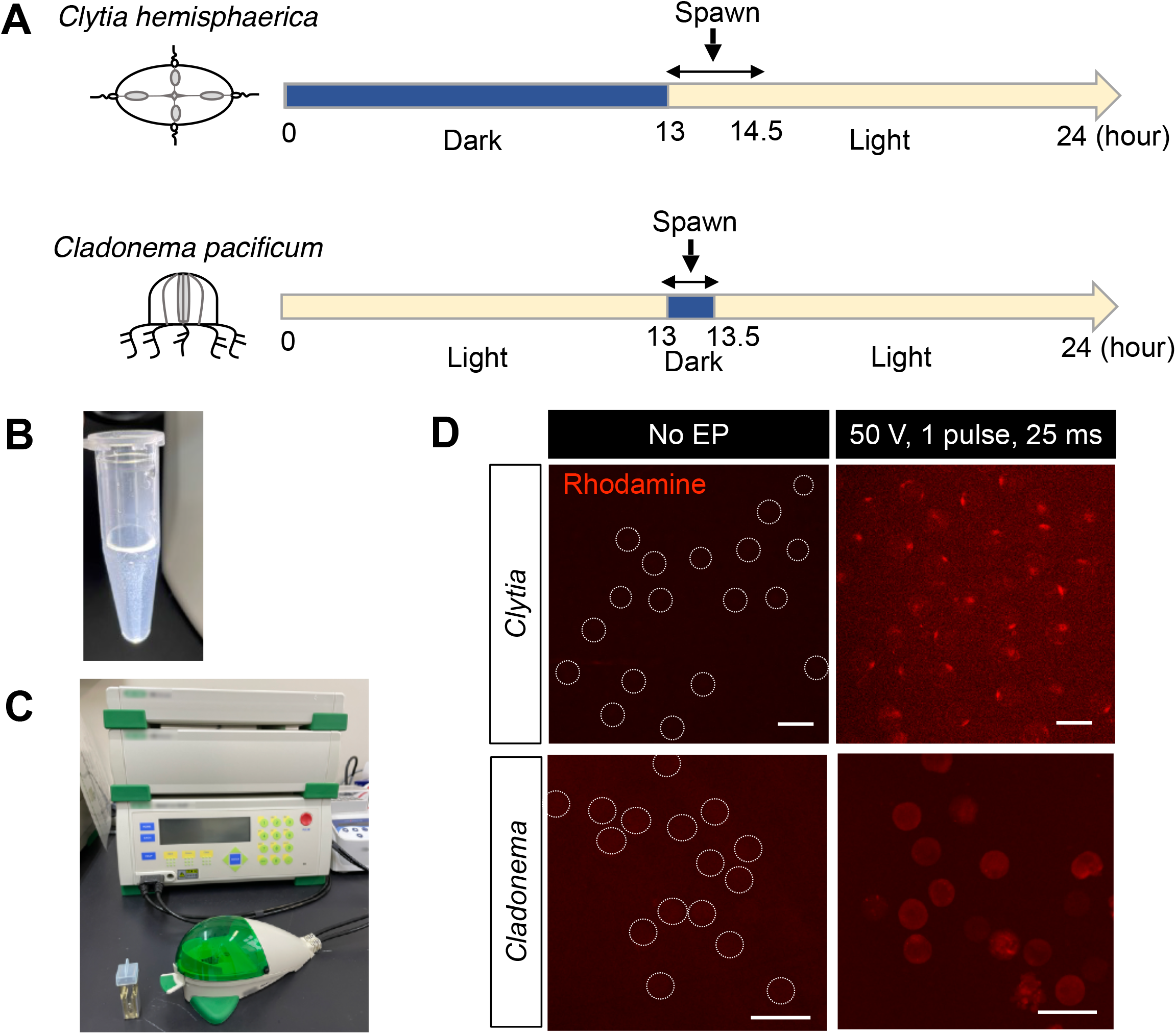
The electroporation procedure for jellyfish eggs. (**A**) Schematic of spawning under the 24 h light/dark cycle. *Clytia hemisphaerica* are maintained on a 13 h dark/11 h light cycle. After light stimuli, sperm spawning occurs within 60-90 min and egg spawning occurs within 90-120 min. *Cladonema pacificum* are maintained on a 23.5 h light/0.5 h dark cycle. After dark stimuli, sperm and egg spawning occurs within 25 min. (**B**) Fertilized eggs are resuspended in 15% Ficoll/artificial sea water to prevent precipitation of eggs during electroporation. (**C**) Electroporation is performed with the Bio-Rad Gene Pulser Xcell electroporation system and a cuvette with a 4 mm gap. (**D**) Visualization of the Rhodamine-Dextran delivery into eggs by electroporation. Rhodamine is shown in red. While little or no fluorescence was observed in eggs without electroporation (No EP), under the condition of a single 50 V pulse for 25 ms, rhodamine signals were detected in both *Clytia and Cladonema* eggs without cell damage. For clarity, egg outlines are indicated with dashed lines for eggs without electroporation. Scale bars: 200 μm.

**Figure 3.**
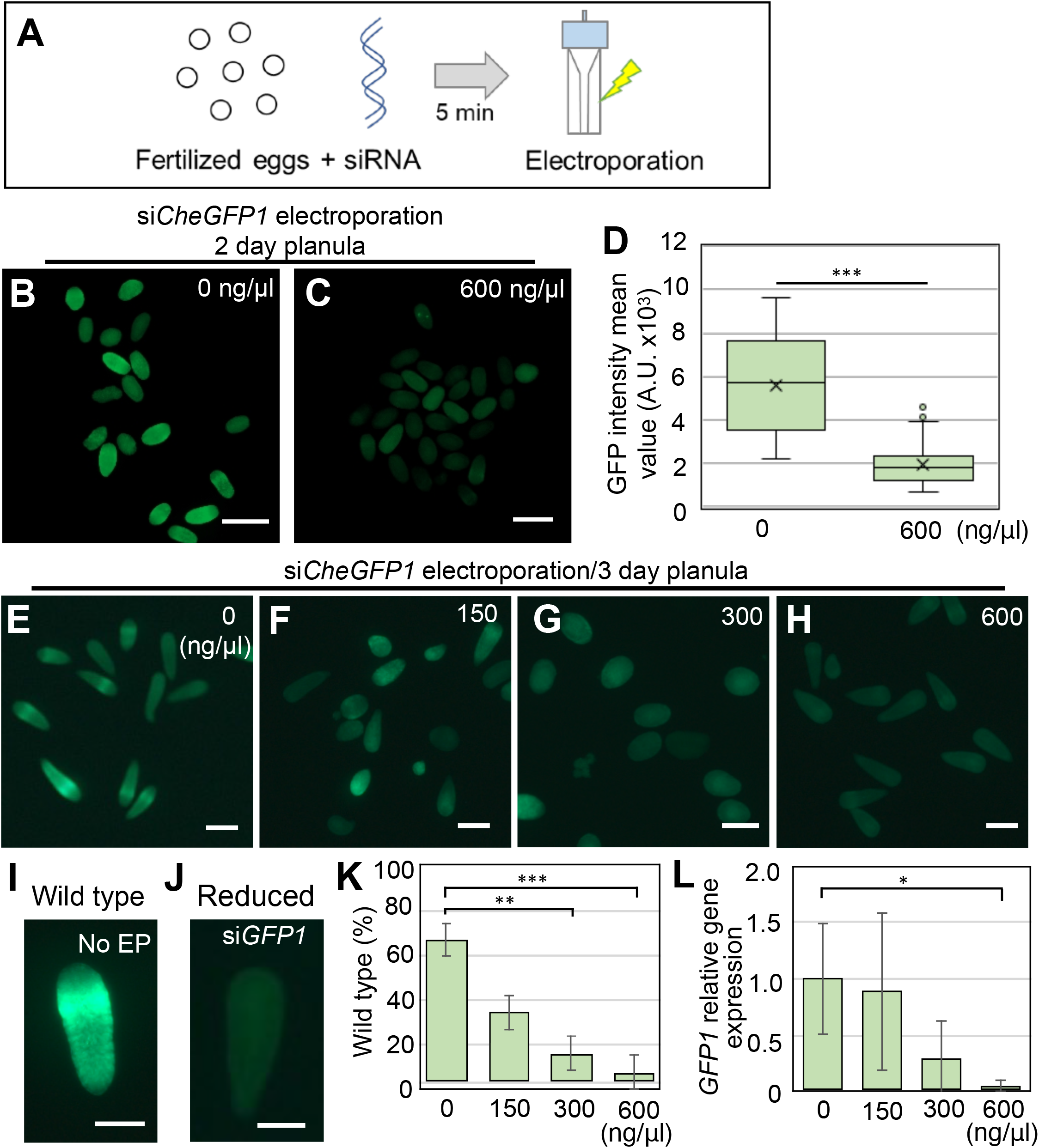
Phenotypes of *GFP1* knockdown with siRNA in *Clytia* fertilized eggs. (**A**) Schematic of the siRNA electroporation procedure using fertilized eggs. Fertilized eggs in 15% Ficoll/artificial sea water mixed with siRNA are incubated for 5 min, and electroporation is conducted after transferring into a cuvette. (**B** and **C**) Phenotypes of *Clytia* 2-day planula after *CheGFP1* siRNA knockdown. (**D**) Boxplots showing the GFP fluorescence intensity mean values in 2-day planula after electroporation of 0 and 600 ng/μl siRNAs targeting *CheGFP1*. Center lines indicate the medians; x’s denote the means; box limits represent the 25th and 75th percentiles; whiskers show the maximum and minimum values. 0 ng/μl, n=24; 600 ng/μl, n=81. ***p<0.001. (**E**-**H**) Phenotypes of 3-day planula after siRNA targeting *CheGFP1* electroporation (0, 150, 300, and 600 ng/μl *siCheGFP1*). (**I** and **J**) For quantification of green fluorescence, we classified phenotypes into two categories: planula with ring-like green fluorescence as “wild-type,” and planula with reduced green fluorescence as “reduced”. (**K**) Quantification of planula larvae positive for *CheGFP1* (wild-type). Bar plots show the percentage of wild-type phenotype after *CheGFP1* knockdown. Number of examined planula: 0 ng/μl, *n*=82; 150 ng/μl, *n*=64; 300 ng/μl, *n*=58; 600 ng/μl, *n*=114. Error bars: maximum and minimum values. Experiments were repeated three times. p-values: 150 ng/μl, p=0.064505; 300 ng/μl, p=0.001437; 600 ng/μl, p=0.00066. **p<0.01, ***p<0.001. (**L**) Quantification of *CheGFP1* mRNA expression levels in 3-day planula by RT-qPCR. *CheEF1alpha* was used as an internal control. Bar heights represent mean values of at least three independent experiments. *CheGFP1* expression levels are standardized relative to the control (0 ng/μl) condition. Error bars: standard deviation. Experiments were performed in triplicated and repeated at least three times. p-values: 150 ng/μl, p=0.829361; 300 ng/μl, p=0.08588; 600 ng/μl, p=0.037359. *p<0.05. Scale bars: 200 μm.

Unfertilized *Clytia* and *Cladonema* eggs were collected and resuspended in 15% Ficoll (Nacalai tesque, Japan) in ASW to prevent the eggs settling from at the bottom of the microtube or sticking to the microtube surface, keeping the egg solution homogeneous (Fig. 2B)^36,38^. Sperm water was added into the egg solution, and the eggs and sperm were incubated for 5 min at room temperature (20-22°C) until fertilization occurred. The sperm water was then removed from the egg solution, and fertilized eggs were resuspended in 15% Ficoll in ASW to adjust the total volume needed for the number of electroporations. For each electroporation, 200-400 of *Clytia* fertilized eggs or 200-700 of *Cladonema* fertilized eggs were prepared in 90 μl of 15% Ficoll in ASW. 10 μl of siRNA/shRNA solution that was adjusted to the proper concentration in RNase free water was added to the fertilized egg solution.

The total 100 μl of fertilized egg and RNA mixture was carefully transferred into a 4 mm cuvette using a 200 μl pipet tip and placed into the shockpod cuvette chamber connected to the Gene Pulser Xcell (Bio-Rad). In most experiments, the Gene Pulser Xcell was used for pulsing with a square wave voltage of 50 V and a single pulse duration of 25 ms, except when optimizing electroporation conditions (Supplementary Figs 1-2). After electroporation, fertilized eggs were transferred to a dish, and incubated for 10 min. The 15% Ficoll solution was removed and replaced with new ASW, and the samples were incubated at 20-22°C.

### Collection of egg sacs, dejellying, and electroporation of *Nematostella* fertilized eggs

The collection of egg sacs, dejellying, and electroporation steps were performed as previously described^36^ with several modifications. Briefly, unfertilized egg sacs were incubated in culture media with sperm at 20-25OC for 30 min. The fertilized egg sacs were then dejellied in freshly prepared L-Cysteine solution (0.04 g/ml, Nacalai tesque, in brackish water; pH adjusted by 5N NaOH at 7.5-8.0) on a shaker for 10 min. The dejellied eggs were rinsed with brackish water before they were suspended in Ficoll solution (Sigma-Aldrich, in brackish water) at a final concentration of 15%. The suspended eggs and siRNA (5 μg/μl stock solution, in MilliQ) were transferred to a 4 mm cuvette (Bio-Rad) with a final volume of 200 μl and were gently mixed. The Bio-Rad Gene Pulser Xcell was used for pulsing, with a square wave voltage of 50 V and a pulse duration of 25 ms. The samples were transferred to a 100 mm dish and maintained in brackish water for 3 days.

### Imaging

GFP and Rhodamine fluorescent imaging as well as bright field imaging, including *in situ* hybridization imaging, were taken with an Axio Zoom V16 (ZEISS), and image processing was done with ZEN 3.4 blue edition (ZEISS) and Fiji/ImageJ software^45^. Intensity of mean values of green fluorescence was measured using the average brightness (pixel value) of the pixels. To prevent planula movement, we added 0.5 M sodium azide (final concentration was less than 0.25M).

### Morphological quantification of planula larvae

After *Wnt3* knockdown by siRNA electroporation of *Clytia* or *Cladonema*, we approximated the planula body shape into a two dimensional ellipse, measured both the long and short axes (Figs. 4C and 5D), and calculated the aspect ratio of those values using Fiji/ImageJ software (Figs, 4D, 5E).

**Figure 4.**
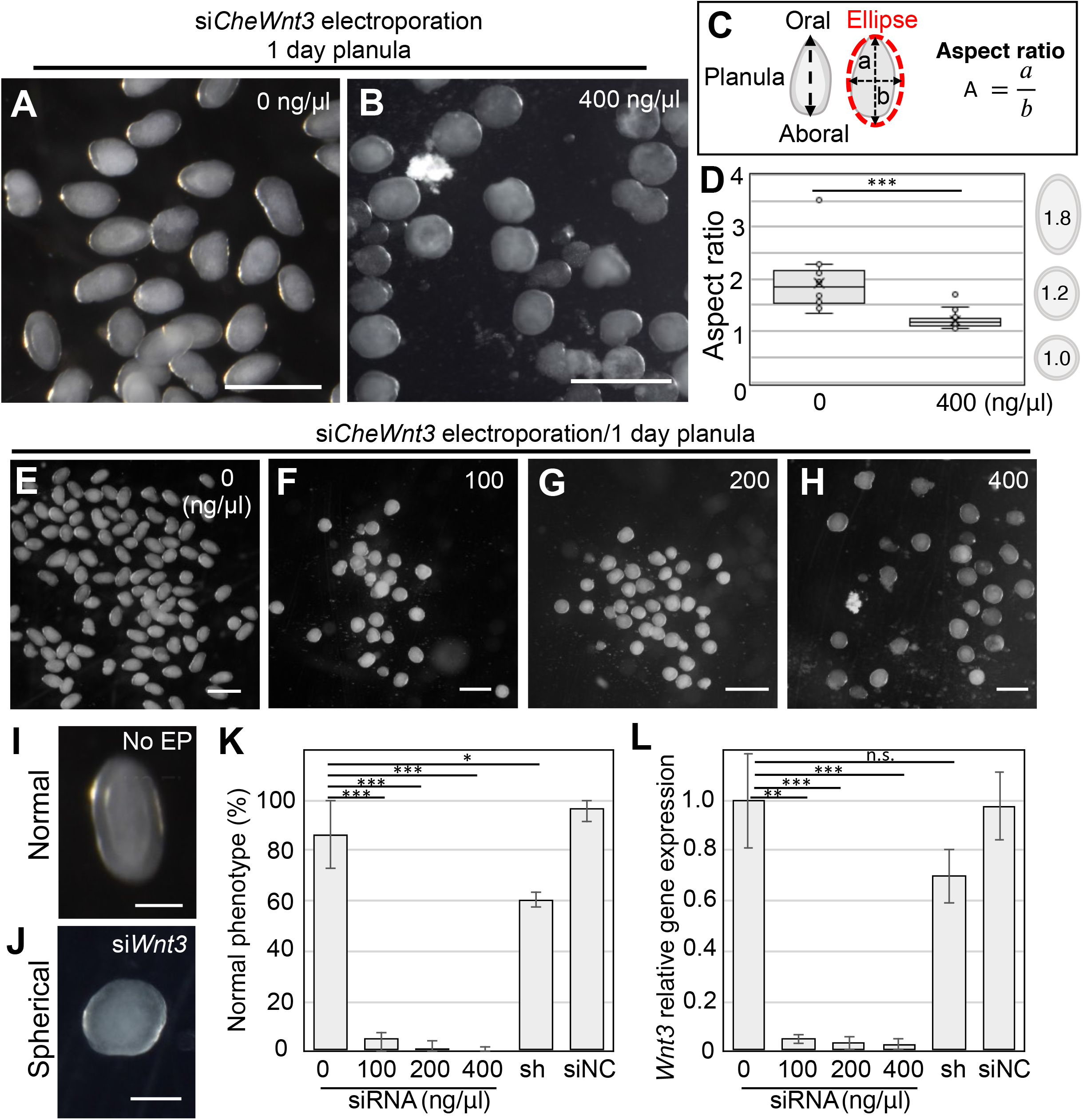
Phenotypes of *Wnt3* knockdown with siRNA in *Clytia* fertilized eggs. (**A** and **B**) Typical phenotypes of *Clytia* 1-day planula. While morphologies of control planula (0 ng/μl) have an elongated oval-shape (A), those of *CheWnt3* siRNA (400 ng/μl) are spherical in shape (**B**). (**C**) Schematic of ellipse approximation for planula morphology and calculation of the aspect ratio. The aspect ratio was calculated by dividing the long axis (a) by short axis (b). (**D**) Boxplots showing the aspect ratio of the 1-day planula. Center lines show the medians; x’s denote the mean values; box limits indicate the 25th and 75th percentiles; whiskers show maximum and minimum values; inner points and outliers are represented by circles. Number of examined planula: 0 ng/μl, *n*=127; 400 ng/μl. *n*=25, ***p<0.001. (**E**-**H**) Phenotypes of 1-day planula after siRNA targeting *CheWnt3* electroporation (0, 100, 200, and 400 ng/μl *siCheWnt3*). (**I** and **J**) For morphology quantification, we classified planula phenotypes into two categories: elongated oval shape as “normal” and spherical (circle) shape as “spherical”. (**K**) Bar plots show the percentage of normal phenotypes across four different siRNA doses as well as planulae treated with shRNA and siNC, a siRNA universal negative control. Percentages are the mean value and error bars indicate standard deviation. Numbers of examined planula: 0 ng/μl, *n*=127; 100 ng/μl, *n*=75; 200 ng/μl, *n*=86; 400 ng/μl, *n*=95. shRNA 400 ng/μl, *n*=288; siNC 400 ng/μl, *n*=45. Experiments were repeated three times. Error bars: maximum and minimum values. p-values: 100 ng/μl, p=0.000599; 200 ng/μl, p=0.000438; 400 ng/μl, p=0.000402; 400 ng/μl of shRNA, p=0.031928; 400 ng/μl of siNC, p=0.283359. *p<0.05, ***p<0.001. (**L**) Quantification of *CheWnt3* mRNA levels of 1-day planula by RT-qPCR. *CheEF1alpha* was used as an internal control. Bar heights represent the mean value. *CheWnt3* expression levels are standardized relative to the control (0 ng/μl). Error bars: standard deviation. p-values: 100 ng/μl, p= 0.00799; 200 ng/μl, p= 0.000754; 400 ng/μl, p= 0.000733; 400 ng/μl of shRNA, p= 0.095403; 400 ng/μl of siNC, p= 0.900264. **p<0.01, ***p<0.001. n.s., not significant. Experiments were performed in triplicate and repeated at least three times. Scale bars: (A, B, E-H) 400 μm, (I, J) 100 μm.

**Figure 5.**
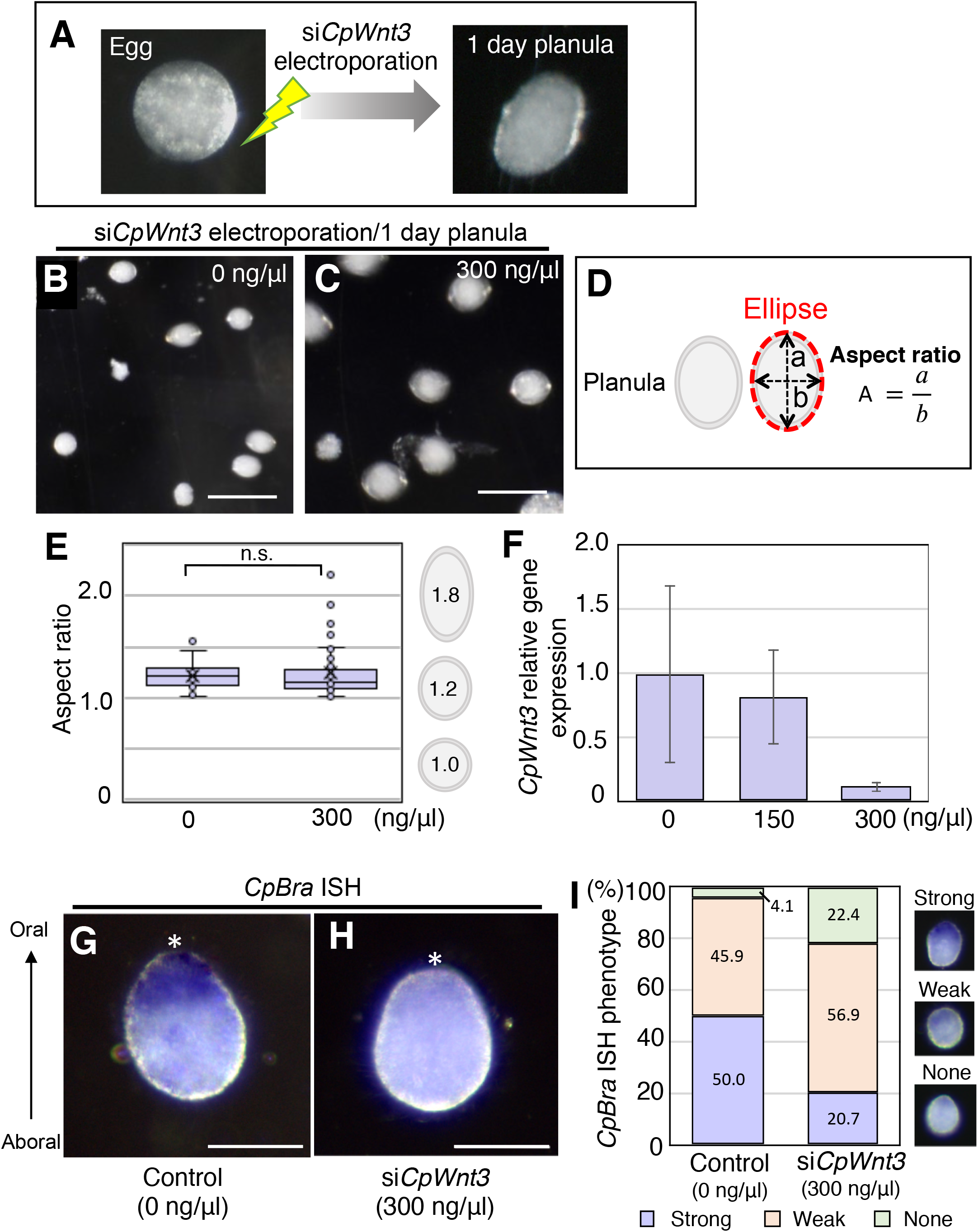
Phenotypes of *Wnt3* knockdown with siRNA in *Cladonema* fertilized eggs. (**A**) Typical morphology of *Cladonema* egg and planula. The wild-type *Cladonema* planula larvae exhibit a slight oval shape. To deliver siRNAs targeting *CpWnt3*, electroporation was performed in fertilized eggs (1-cell stage), and phenotypes were confirmed at planula in one day. (**B** and **C**) Phenotypes of 1-day planula after *CpWnt3* siRNA knockdown. (**D**) Schematic of ellipse approximation for planula morphology and calculation of the aspect ratio. The aspect ratio was calculated by dividing the long axis (a) by short axis (b). (**E**) Boxplots showing the aspect ratio of the 1-day planula after *siCpWnt3* electroporation (0 and 300 ng/μl). Center lines show the medians; x’s denote the mean values; box limits indicate the 25th and 75th percentiles; whiskers show maximum and minimum values; inner points and outliers are represented by circles. Number of examined planulae: 0 ng/μl, *n*=22; 300 ng/μl, *n*=50. p= 0.658303. n.s., not significant. (**F**) Quantification of *CpWnt3* mRNA levels in 1-day planula by RT-qPCR. *CpEFIalpha* was used as an internal control. Bar heights represent the mean value. Error bars indicate standard deviation. *CpWnt3* expression levels are standardized relative to the control (0 ng/μl). Experiments were performed in triplicate and repeated three times. p-values: 150 ng/μl, p= 0.376498; 300 ng/μl, p= 0.073956. (**G, H**) Representative *in situ* hybridization (ISH) image of oral *CpBra* expression in 1-day planula for control (G: 0 ng/μl si*CpWnt3*) and *CpWnt3* siRNA (H: 300 ng/μl si*CpWnt3*). The oral pole is indicated by an asterisk. (**I**) Quantification of *CpBra* expression phenotypes based on ISH images. Stacked bar plots show the percentage of 1-day planulae in each phenotypic class. *CpBra* expression patterns are categorized into three phenotypes (strong, weak and none). 0 ng/μl, *n*=74; 300 ng/μl, *n*=58. Scale bars: (B, C) 200 μm, (G, H) 50 μm.

### RT-qPCR

For *Clytia* and *Cladonema* samples, total RNA was extracted with RNeasy Mini or Micro kits (Qiagen). Lysate was treated with DNaseI (Qiagen) for 15 min at room temperature (RT). cDNA was synthesized with the PrimeScript II 1^st^ strand synthesis kit (Takara bio). RT-qPCR was performed with CFX connect (Bio-Rad) using iTaqTM universal SYBER Green Supermix (Bio-Rad) or the QuantStudio 6 Flex Real-Time PCR System (Thermo Fisher) using TB Green Premix Ex TaqII (Tli RNaseH Plus) (Takara, RR820). Gene expression was normalized to the housekeeping gene, *EF1alpha* or *F-actin capping protein subunit beta*, and the delta-delta-ct method was used for quantification (CFX maestro software, Bio-Rad or QuantStudio 6 Flex Real-Time PCR System software, Thermo Fisher).

For *Nematostella* samples, total RNA was extracted using the RNeasy Mini Kit and RNase-Free DNase Set (Qiagen). cDNA synthesis was conducted using the SuperScript IV First-Strand Synthesis System (Thermo Fisher). RT-qPCR was performed with the StepOnePlusTM Real-Time PCR System (Applied Biosystems, Thermo Fisher) using PowerUP SYBR Green Master Mix (Thermo Fisher). Gene expression was normalized to the housekeeping gene, *NvEf1alpha*, and the delta-delta-ct method was used for quantification (StepOne™ and StepOnePlus™ Software v2.3).

### *In situ* hybridization

Purified total RNA was reverse transcribed into cDNA by PrimeScript™ 2 1st strand cDNA Synthesis Kit (TaKaRa). The target gene fragments *(Brachyury, CpBra)* were amplified from a cDNA library. The primer sets used for PCR cloning are as follows: *CpBra:* 5’-GCTCCCATAAGATCCGGTCG −3’(forward) and 5’-TTTGTCGCAGTCGAAGACCA-3’(reverse), or 5’-GAACGGTGATGGACAGGTCA-3’ (forward) and 5’-GGATTCCAAGGATTGGGCGT-3’ (reverse). PCR products were sub-cloned into the TAK101 vector (TOYOBO). The resulting plasmids were used for RNA probe synthesis with digoxigenin (DIG) labeling mix (Roche), and T7 or T3 RNA polymerase (Roche) was used, according to the insert direction. The detail for the probe synthesis was referred to the published protocol^46^.

Planula larvae were anesthetized in 50% 0.5 M NaN3 in ASW for 5 min and fixed overnight at 4°C with 4% paraformaldehyde (PFA) in ASW. The following whole mount *in situ* hybridization was performed following the protocol in Hou et al.^24^ with slight modifications. Briefly, fixed samples were washed three times with PBS containing 0.1% Tween-20 (PBST), followed by pre-hybridization in hybridization buffer (HB buffer: 5×SSC, 50% formamide, 0.1% Tween-20, 50 μg/ml tRNA, 50 μg/ml heparin) at 55°C for 2 h. Samples were hybridized with HB Buffer containing the antisense probes (final probe concentration: 0.5-1 ng/μL in HB Buffer) at 55°C overnight. The samples were washed each twice with wash buffer 1 (5x SSC, 50% formamide, 0.1% Tween-20), wash buffer 2 (2x SSC, 50% formamide, 0.1% Tween-20) and 2x SSC. The samples were then washed with 0.1% PBST and incubated in 1% blocking buffer (1% blocking reagent [Roche, 11175041910] in Maleic acid) for 1 h. The samples were incubated with alkaline phosphatase (AP)-conjugated anti-DIG antibodies (1:2000, Roche, 11175041910) in 1% blocking buffer for 4 h at room temperature. Colorimetric reactions were performed by NBT/BCIP (Roche, 11175041910) in alkaline phosphatase buffer (100 mM NaCl, 100 mM Tris-HCl (pH 9.5), 50 mM MgCl2, 0.1% Tween-20) until the signals were detected.

### Sequence alignment and phylogenetic analysis

Amino acid sequence alignment of Wnt3 and Wnt family proteins were performed by ClustalW^47^. GenBank accession numbers and protein names are listed in Supplementary Table 3 and 4. The alignment data was visualized on Jalview software^48^. Maximum likelihood (ML) tree calculation was inferred on ClustalW with the PhyML bootstrap method. Branch supports were computed out of 100 bootstrapped trees. The tree visualization was created using MEGA-X software^49^.

### qPCR primers

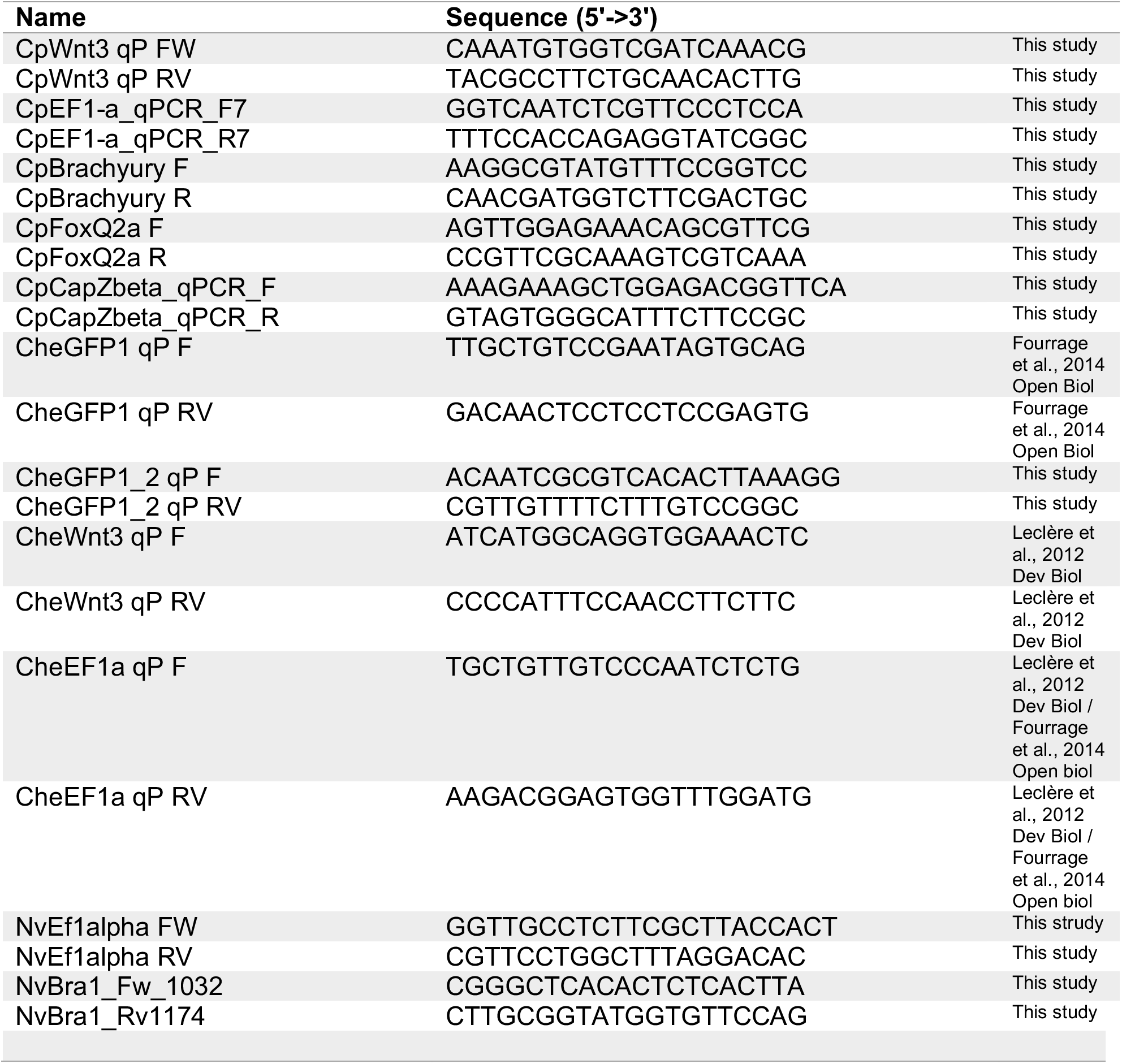

### siRNA design

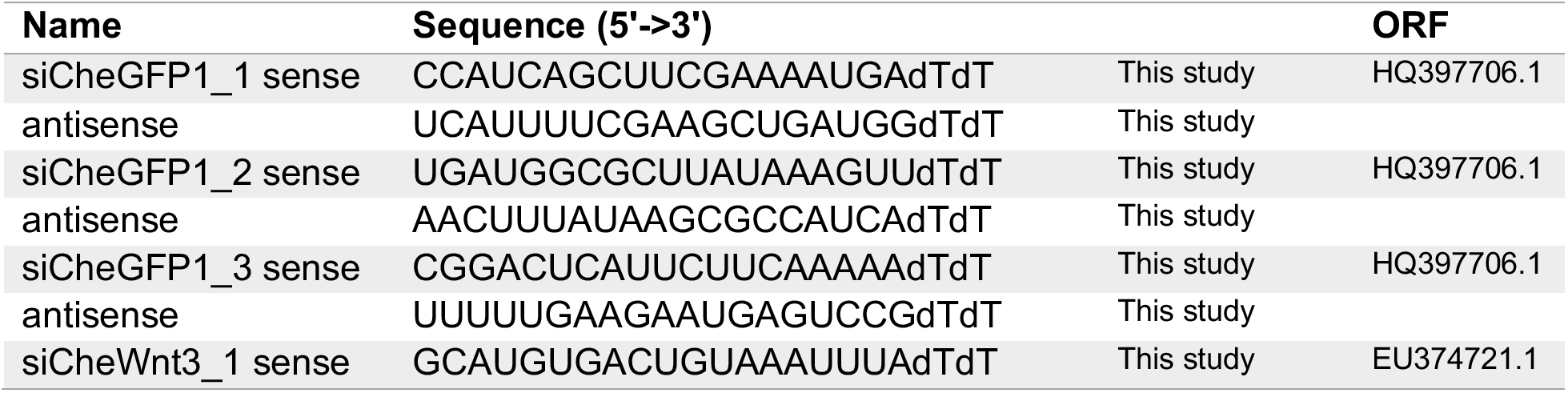
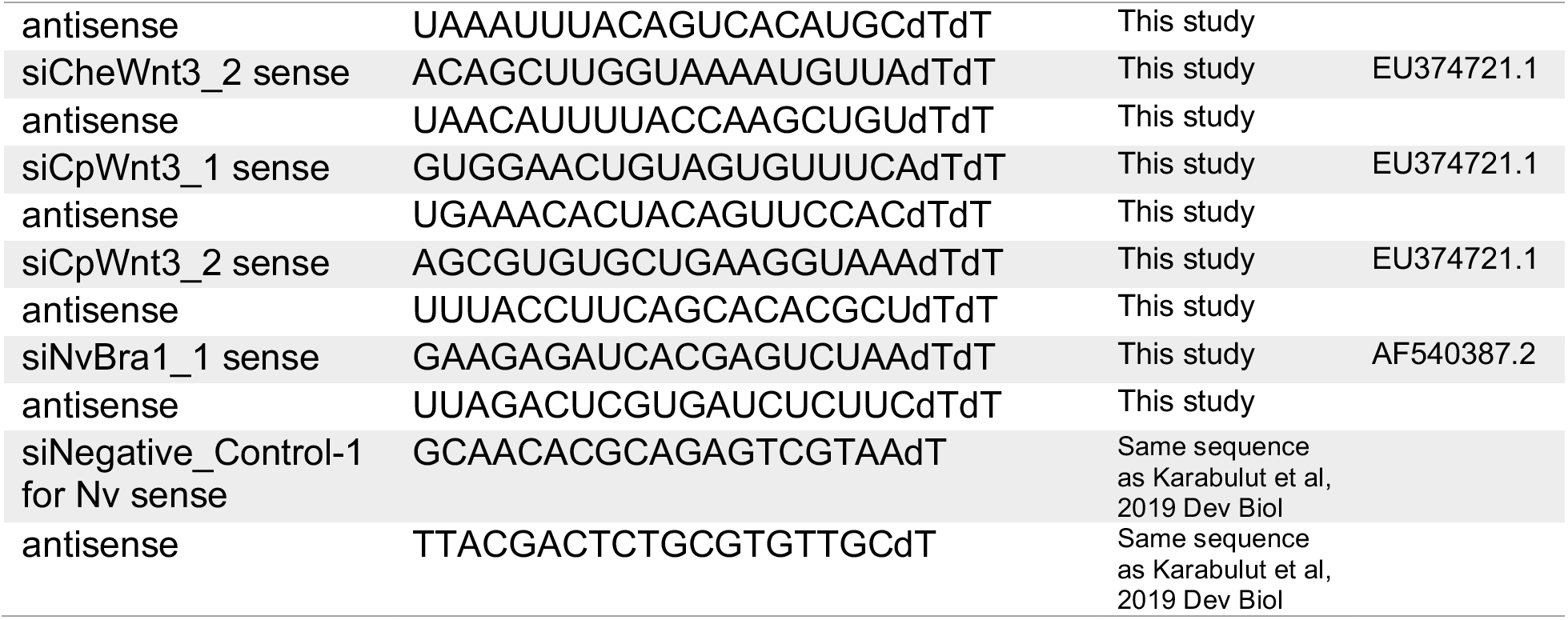

### shRNA template

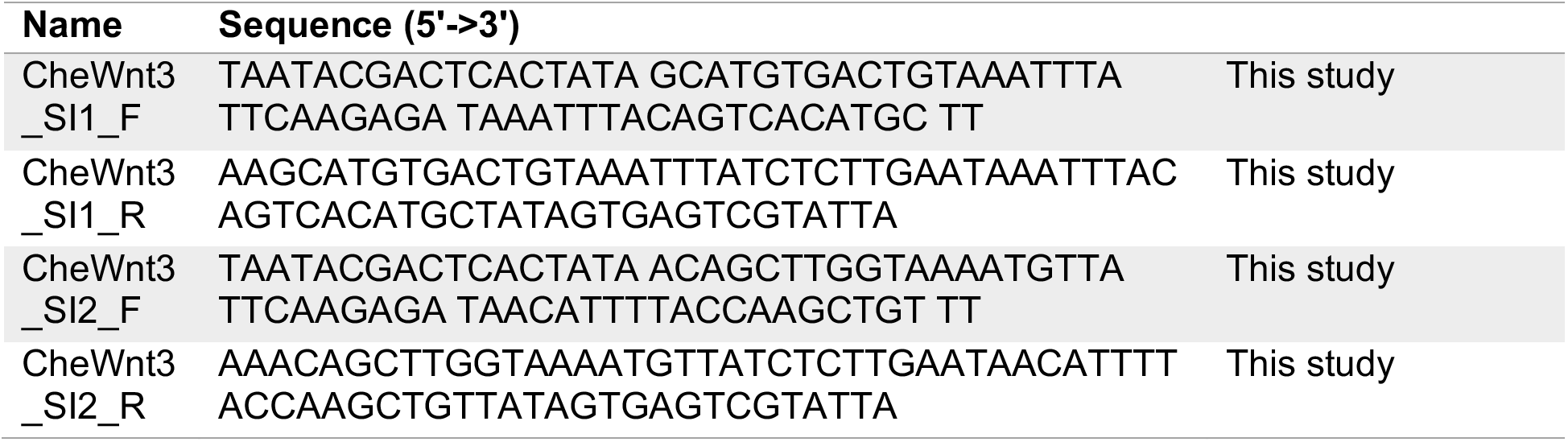

### Graphs and statistical analysis

All graphs were prepared in Excel. To assess phenotypes and RT-qPCR statistical significance, we used the percentage values and delta-delta-Ct values to perform two-tailed Student’s t tests, except in Figure 5F, where a one-tailed Student’s t test was used. For statistics related to fluorescence intensity mean value and aspect ratio, Kolmogorov–Smirnov normality tests were used. All statistical tests were performed at https://www.socscistatistics.com.

## Results

### Optimization of electroporation conditions

In both hydrozoan jellyfish *Clytia hemisphaerica* and *Cladonema pacificum*, gametogenesis is regulated by light-dark transitions^6,50^. A previous report showed that *Clytia* medusae release eggs within 110-120 min and sperm within 60-90 min upon light illumination after 3-8 h of darkness^12^. *Cladonema pacificum*, which live along coastal areas in Japan, exhibit two types of gametogenesis depending on habitat: the dark-light transition (light stimulation type) and the light-dark transition (dark stimulation type), respectively. Dark stimulation *Cladonema* medusae, which we used in this study, release eggs or sperm within 30 min of dark stimulus after 20-24 h of constant exposure to light^25,50^. By utilizing the characteristic gametogenesis of these jellyfish species, we can control egg/sperm release to induce fertilization at any time of day. Indeed, we routinely obtained *Clytia* sperm within 60 min and *Clytia* eggs within 90 min of light on a 13h dark/11h light cycle (Fig. 2A). Similarly, we collected gametes from *Cladonema* within 25 min of dark stimulation on a 23.5h light/30 min dark cycle (Fig. 2A).

To determine optimal electroporation parameters, we first tested different electroporation conditions and monitored the delivery efficiency of the red fluorescent dye Rhodamine-Dextran into unfertilized *Clytia* and *Cladonema* eggs (Supplementary Figs. 1-2). Given that the average molecular weight of siRNA (21-mer, ~13,300 Da) is similar to the molecular size of Rhodamine-Dextran (10,000 Da), electroporation trials with Rhodamine-Dextran can visually confirm that small molecules are incorporated into jellyfish eggs, as previously shown in *Hydractinia* eggs^38^. The collected eggs were suspended in 15% Ficoll artificial seawater to prevent precipitation and to make a homogeneous solution (Fig. 2B)^36,38^. We performed electroporation trails for Rhodamine-Dextran using a cuvette with a 4 mm gap and tested several conditions of voltage (V) and pulse length time (milliseconds, ms) using a conventional electroporation system (Fig. 2C). As previous work showed that increasing the number of pulses does not increase the Dextran delivery rate but decreases the survival rate of embryos^38^, we fixed the number of pulses to one and incubated unfertilized eggs with 1 mg/ml Dextrane-Rhodamine solution. *Clytia* eggs have little autofluorescence in the red channel, and without electroporation (No EP), red fluorescence was rarely detected in unfertilized eggs (Fig. 2D and Supplementary Fig. 1A). We compared the eight different electroporation conditions and found that the rhodamine fluorescence was detected in *Clytia* eggs and correlated with the strength of electric voltage, as long as the voltage was in the range of 50-100 V (Supplementary Figs. 1B-1D). By contrast, when the voltage exceeded 200 V, the eggs ruptured from physical damage, and cellular debris was observed (Supplementary Figs, 1E-1G).

We performed similar electroporation trials on unfertilized *Cladonema* eggs under eight different conditions. While red autofluorescence was negligible in the *Cladonema* eggs in the absence of electroporation (Fig. 2D and Supplementary Fig. 2A), Rhodamine fluorescence was frequently detected with 50 V (Fig. 2D and Supplementally Fig. 2E). In the more intense conditions beyond 100 V (Supplementally Figs. 2F and 2G), the eggs were severely damaged. Based on these results, we established the optimal electroporation conditions (50 V, 1 pulse, 25 ms) that provide high frequency of Dextran-positive eggs and low levels of cell damage for both jellyfish eggs (Fig. 2D). Of note, during the initial trials, we used unfertilized eggs, but we found that the electroporation process severely affected the survival rate of subsequent embryos. In the case of *Cladonema*, electroporation before fertilization severely decreased the survival rate of fertilized eggs (7.04% at 1 h post fertilization) while electroporation after fertilization did not influence the survival rate (No EP: 59.67%, With EP: 60.15%) (Supplementary Table 1). This dissimilarity is likely due to the decrease of fertilization rate after electroporation, and both elapsed time after spawning and physical damage by electroporation can affect fertilization efficiency. Therefore, we decided to conduct further electroporation experiments with fertilized eggs.

### Gene knockdown with siRNA in *Nematostella vectensis*

Before attempting gene knockdown via siRNAs electroporation in fertilized jellyfish eggs, we first used *Nematostella vectensis*, where egg electroporation methodology has been established^36^. We selected *Nematostella Brachyury* (*NvBra*), a gene expressed in the blastopore margin that functions in the early embryo^51,52^, as a target. After electroporation of siRNA targeting *NvBra* in fertilized eggs, we examined *NvBra* expression in planula larvae at 3 days-post-fertilization (dpf) by RT-qPCR and found that *NvBra* expression was drastically suppressed compared to no electroporation controls and negative control siRNAs (Supplementary Fig. 3). We further evaluated survival rates after electroporation and confirmed that siRNA electroporation does not severely affect survival rates of 3 dpf planula in any of the tested conditions (Supplementary Table 2). These results suggest that siRNA delivery into fertilized eggs via electroporation is an effective gene knockdown method that is comparable to the effect of shRNA delivery into unfertilized eggs via electroporation^36^, supporting our attempt to perform gene knockdown via siRNA electroporation using fertilized jellyfish eggs.

### Endogenous *GFP1* knockdown with siRNA in *Clytia hemisphaerica*

The hydrozoan jellyfish *Clytia* endogenously expresses green fluorescent proteins, which are encoded by 14 genes grouped into 4 clusters *(CheGFP1-4)*^16^. The expression pattern for each GFP changes by stage in the life cycle: for instance, *Clytia* eggs express maternal *CheGFP2* mRNA, and planula larvae show typical ring-like *CheGFP1* fluorescence in the lateral ectoderm at 3 dpf^11^. While *CheGFP1* gene expression becomes dominant in the planula stage; *CheGFP2, CheGFP3*, and *CheGFP4* are expressed in different tissues and organs in the medusa^53^. Momose *et al.* succeeded in genome editing by injecting CRISPR/Cas9 targeting the endogenous *CheGFP1* into fertilized eggs, and confirmed the knockout effect by loss of green fluorescence in 3 dpf planula larvae^11^. In addition, *CheGFP1* is not essential for animal survival, which allows for the monitoring of knockdown effects after embryogenesis. For these reasons, we chose endogenous *CheGFP1* for the first target gene of siRNA knockdown.

In order to maximize the effect of gene knockdown, we used a combination of three different siRNAs against *CheGFP1* (Accession No. HQ397706.1; Supplementary Fig. 4A) in a mixture with a 1:1:1 ratio, and electroporated 600 ng/μl of siRNA mix (200 ng/μl per each siRNA) into fertilized eggs (Fig. 3A). To verify the effect of gene knockdown, we measured green fluorescence intensity in 2 dpf planula and found that siRNA (600ng/μl siRNA mix) electroporated embryos showed dramatically reduced fluorescence intensity compared to the no-siRNA controls (Figs. 3B-D; mean intensity value: 5.7×10^3^ for 0 ng/μl control, 1.8×10^3^ for 600 ng/μl). To evaluate the knockdown effects at varying doses of siRNA, we next electroporated different concentrations of the *CheGFP1* siRNA mix (0, 150, 300, and 600 ng/μl) into fertilized eggs. While control 3 dpf planulae with no-electroporation (No EP) as well as those with no-siRNA showed typical ring-shape GFP fluorescence in the ectoderm, many planulae with *CheGFP1* siRNA electroporation lost green fluorescence (Figs. 3E-3H). To evaluate the GFP knockdown effects, we classified planulae with typical GFP1 fluorescence as “wild type” and planulae with reduced fluorescence as “reduced” for further quantification (Figs. 3I and 3J). The percentage of wild type phenotype dramatically reduced in an siRNA dose-dependent manner (Fig. 3K). To further quantify the above result on a molecular level, we examined relative *CheGFP1* mRNA levels by RT-qPCR using the primer set that was evaluated in a previously study^53^. Consistent with the image analysis results, we confirmed a marked reduction of *CheGFP1* gene expression at different concentrations of *CheGFP1* siRNA (Fig. 3L). These results demonstrate that siRNAs targeting *CheGFP1* repress its expression upon electroporation and that the knockdown effects are dose-dependent.

Given the presence of multiple *CheGFP1* loci (seven nearly identical *CheGFP1* genes) in the *Clytia* genome^16^, it was not clear whether three siRNAs against *GFP1* sufficiently suppress gene expression in different versions of *CheGFP1*, particularly those expressed in the planula stage (Supplementary Fig. 4A). We thus performed RT-qPCR using the newly designed primer set that amplifies all seven *CheGFP1* genes and found a dramatic reduction of gene expression upon electroporation of *CheGFP1* siRNAs (600 ng/μl) (Supplementary Fig. 4B). Altogether, these results indicate that our combination of *CheGFP1* siRNAs suppresses gene expression of *CheGFP1* in the *Clytia* planula.

### *Wnt3* knockdown with siRNA in *Clytia hemisphaerica*

Since electroporation of siRNAs targeting endogenous *CheGFP1* was shown to be effective, we next decided to investigate the effects of knocking down other genes that play important roles in early development. Wnt3 is a secreted signaling protein in the Wnt/β-catenin pathway that is involved in diverse developmental processes including cell fate decisions and patterning^54,55^. In *Clytia, Wnt3* RNA is maternally localized to the animal cortex, the future oral side of the embryo, and Wnt3 organizes axial patterning as the main ligand for the Wnt/β-catenin pathway in the early embryonic stage^10^. After embryogenesis, the *Clytia* planula normally elongates along the oral-aboral axis and becomes oval in shape (Fig. 4A). By contrast, after *CheWnt3* morpholino injection, the morphant loses oral-aboral axis polarity and exhibits a spherical shape^10^. This raises the possibility that knockdown of *CheWnt3* via siRNA may result in a similar spherical morphology for *Clytia* larvae as is observed in *CheWnt3* morphants. Therefore, to evaluate siRNA knockdown effects on *Clytia*, we chose *CheWnt3* as our second target.

We used a mixture of two different siRNAs (1:1 ratio) targeting *CheWnt3*, and carried out electroporation of the siRNA mix into *Clytia* fertilized eggs. After electroporation of *CheWnt3* siRNAs, 1 dpf planula larvae showed nearly complete spherical morphology (Fig. 4B), which is reminiscent of *CheWnt3* morphants^10^. To quantify the effect of siRNA knockdown on *Clytia* larval morphology, we calculated the aspect ratio in two dimensions, where values greater than 1.0 shows the tendency to be oval (ellipsoid-like) while a value of 1.0 indicates a perfect circle (sphere) (Fig. 4C). The control larvae with 0 ng/μl siRNA showed a median aspect ratio of 1.69, indicating a typical ellipsoid-like morphology. By contrast, larvae that underwent electroporation with *CheWnt3* siRNAs (400 ng/μl) showed a median aspect ratio of 1.16, suggesting a much greater tendency toward spherical morphology (Fig. 4D). These results quantitively confirmed the effect of *CheWnt3* knockdown on larval morphology, which phenocopies *CheWnt3* morphants^10^. To analyze the level of gene-specific knockdown at varying doses of siRNAs, we next electroporated mixtures of 0, 100, 200, and 400 ng/μl of *CheWnt3* siRNAs into *Clytia* fertilized eggs (Figs. 4E-4H). To evaluate *CheWnt3* siRNA effects on morphology, we classified 1 dpf planulae with elongated ellipsoid-like shapes as “normal” and planulae with spherical shapes as “spherical” (Figs. 4I and 4J). The percentage of normal phenotype dramatically reduced after *CheWnt3* siRNA-electroporation (< 1.0 % in 400 ng/μl), even at the lowest dose (5.2% in 100 ng/μl) (Fig. 4K). To further evaluate the result at a molecular level, we analyzed relative *CheWnt3* mRNA levels by RT-qPCR and confirmed a significant reduction in *CheWnt3* gene expression at all concentrations (Fig. 4L). Of note, after electroporation with universal negative control siRNA (Nippon Gene Co., Ltd.), both morphological phenotypes and relative gene expression were comparable to no-siRNA controls (Figs. 4K and 4L), further confirming that siRNA-mediated knockdown by electroporation is target-specific. These results together indicate that siRNAs targeting endogenous genes effectively knock down their expression in *Clytia* embryos.

Effective gene knockdown via shRNA electroporation has previously been reported using two cnidarian polyps, *Nematostella vectensis* and *Hydractinia symbiolongicarpus*^36,38^. To test whether shRNAs are also effective in the hydrozoan jellyfish *Clytia*, we designed shRNAs targeting *CheWnt3* using the same 19 bp target sequence that was used for siRNAs, and performed electroporation on fertilized eggs with a mix of two different *Wnt3* shRNAs. After electroporation of *Wnt3* shRNAs, relative *Wnt3* mRNA expression in 1 dpf planula was weakly down-regulated (0.83), and was not significant compared to the striking effects of *Wnt3* siRNA electroporation (0.039) at the same concentration (400 ng/μl) (Fig. 4L). In addition, electroporation of *Wnt3* siRNAs resulted in an extensive morphological change toward spherical shaped planulae, showing almost no “normal” phenotype (0.71%), whereas electroporation of shRNA had a limited effect (60.4%) (Fig. 4K). These results suggest that the effect of siRNA is much stronger than that of shRNA in *Clytia* when treated with the same target sequence and concentration.

### Identification of *Wnt3* in *Cladonema pacificum*

In contrast to *Clytia*, the sole genetic model jellyfish, previous studies have neither established genetic manipulation nor performed genome sequencing of *Cladonema pacificum*, despite its unique biological features. This is partly because the extremely small size of the *Cladonema* egg makes microinjection difficult. By testing whether siRNA-mediated knockdown via electroporation can work in *Cladonema*, we can start to manipulate genes in *Cladonema* while also demonstrating that siRNA electroporation can be applicable to other jellyfish species. While green fluorescent protein (GFP) is often an easy early target for gene manipulation in emerging organisms, *Cladonema* does not exhibit apparent endogenous green fluorescence expression at any stage, unlike *Clytia*. We instead turned our focus to *Wnt3* as the target for gene knockdown in *Cladonema* since Wnt3 plays an important role in early embryonic development in several cnidarian species^56,57^.

To identify the *Wnt3* ortholog in *Cladonema*, we utilized RNA-seq results from the *Cladonema* polyp, stolon, and medusa manubrium (data not shown). We performed *Cladonema Wnt3* CDS annotation using the *Clytia* CDS database (MARIMBA, Marine models database: http://marimba.obs-vlfr.fr) and found one contig annotated with *Wnt3 (CpWnt3)* and multiple *Wnt* genes (Supplementary Figs. 5 and 6). We then performed phylogenic analysis using the neighbor-joining method, and confirmed *CpWnt3* (LC720435) is grouped with medusozoan *Hydra vulgaris (HvWnt3)* and *Clytia Wnt3 (CheWnt3)* rather than Anthozoa *Nematostella* and *Acropora digitifera* (Supplementary Fig. 5A). Multiple sequence alignment further showed highly conserved amino acids sequences among different species including bilaterians and cnidarians (Supplementary Fig. 5B). These findings raise the possibility that *CpWnt3* has a similar function in controlling the Wnt/β-catenin pathway in *Cladonema*.

### *Wnt3* knockdown with siRNA in *Cladonema pacificum*

Does *Wnt3* function during *Cladonema* embryogenesis, particularly during axial patterning? Interestingly, *Cladonema* planula larvae do not show a clear elongated shape (Fig. 5A), as observed in *Clytia*. It is thus possible that morphogenesis, including axial patterning and/or Wnt/β-catenin pathway function, differs between these two jellyfish species. It is also possible that the limited elongation in *Cladonema* planulae might hamper phenotypical appearance upon inhibition of Wnt/β-catenin signaling.

In order to verify that siRNA electroporation works in *Cladonema* and that *Wnt3* affects axial patterning, we performed electroporation of siRNAs targeting *CpWnt3*. We prepared fertilized *Cladonema* eggs as we did in *Clytia*, and used a mixture of two different siRNAs against *CpWnt3* CDS with a 1:1 ratio (300 ng/μl) and performed electroporation with the previously established parameters (50V, 1 pulse, 25 ms) in fertilized *Cladonema* eggs. After electroporation of siRNAs for *CpWnt3*, planula larvae did not exhibit morphological differences compared to controls (Figs. 5A-5C; median aspect ratio: 1.21 for 0 ng/μl control, 1.16 for 300 ng/μl for si*CpWnt3*). We then quantified the morphological phenotypes by calculating the aspect ratio of 1 dpf planulae and confirmed that *CpWnt3* knockdown does not cause a significant morphological change (Figs. 5D and 5E), which is consistent with the possibility that *Cladonema* planulae simply exhibit limited elongation.

Another possibility that would explain the above result is a defect in the siRNA knockdown itself. To test this potential explanation, we carried out RT-qPCR using mRNA samples from 1 dpf planula after electroporating different concentrations of *CpWnt3* siRNA mix (0, 150, and 300 ng/μl) into fertilized eggs. We confirmed a reduction of *Wnt3* gene expression in an siRNA dose-dependent manner (Fig. 5F), eliminating siRNA knockdown defects as the cause of our initial result.

To further confirm whether the Wnt/β-catenin pathway is affected by *Wnt3* knockdown, we examined the gene expression of axial markers *Brachyury (CpBra)* and *FoxQ2a (CpFoxQ2a)*, whose expression are influenced by the Wnt/β-catenin pathway in *Clytia*^10^. From RT-qPCR, we found that *CpWnt3* knockdown causes a decrease in *CpBra* gene expression and an increase in *CpFoxQ2a* gene expression (Supplementary Fig. 7). We also examined *CpBra* expression by *in situ* hybridization and confirmed the reduction of *CpBra* expression on the oral side of planula larvae (Figs. 5G-5I), which is similar to the phenotype exhibited by *CheWnt3* morphants in *Clytia*^10^. Notably, after *CpWnt3* siRNAs electroporation, the rate of metamorphosis from planula to primary polyp decreased dramatically (Supplementary Table 5), implying potential developmental defects upon abrogation of the Wnt/β-catenin pathway. These results suggest that, although overall oral-aboral polarity is less prominent in *Cladonema* compared to *Clytia* at the morphological level, the axial patterning mechanism mediated by Wnt/β-catenin signaling may be conserved between these two jellyfish species.

## Discussion

In this study, we have established a method to knock down endogenous genes in two hydrozoan jellyfish, *Clytia hemisphaerica* and *Cladonema pacificum*, via the siRNA electroporation of fertilized eggs. We showed that knockdown of endogenous *GFP1* in *Clytia* causes the loss of GFP fluorescence in the planula stage (Fig. 3), as previously achieved by CRISPR/Cas9-mediated *GFP1* knockout^11^. We also confirmed that knockdown of *Wnt3* in *Clytia* induces spherical morphology of the planula (Fig. 4), mirroring the results of injections of *Wnt3* morpholino antisense oligo^10^. We further succeeded in efficient repression of *Wnt3* gene expression in *Cladonema* after *Wnt3* siRNA electroporation and found that expression of axial patterning genes is controlled by the Wnt/β-catenin pathway (Fig. 5), suggesting that the conserved mechanism is involved in embryogenesis and planula morphogenesis across hydrozoan jellyfish species.

Our results show that the knockdown efficiency of siRNA is much greater than that of shRNA for electroporation of *Clytia* fertilized eggs with the same *Wnt3* target sequence in the same concentration (Fig. 4K-4L). One possibility that would explain such distinct effects between siRNA and shRNA electroporation is the existence of differences in the RNAi machinery or RNA processing efficiency among cnidarians. During RNAi in mammals, the RNase III Dicer protein processes shRNA in collaboration with cofactors TRBP (Transactivation response element RNA-binding protein) and PACT(protein activator of the interferon-induced protein kinase)to produce a mature form of siRNA^39^. Mature siRNA in the Dicer and TRBP/PACT complex is associated with the Argonaute protein and cleaves endogenous complementary mRNAs as the RNA-induced silencing complex ^30,58^. These small RNA biogenesis factors, Dicers and cofactors (TRBP in bilaterians, HYL1 in plants and cnidarians), are found in *Nematostella vectensis, Acropora digitifera*, and *Hydra vulgaris*^41^. Although Dicers (TCONS_00004571 and TCONS_00004525) and Hyl1 (TCONS_00010722) are predicted to exist in *Clytia* based on MARIMBA transcriptome data, the expression level of Dicers in oocytes and early gastrulation stages is lower than that in other stages, suggesting the possibility that insufficient shRNA to siRNA processing efficiency is responsible for the difference in knockdown efficiency between siRNA and shRNA electroporation. It will be interesting to elucidate the detailed molecular mechanism of RNAi in different cnidarian species and address RNA processing efficiency in distinct developmental stages. The lower efficiency of knockdown by shRNA compared to siRNA in *Clytia* could also be simply explained by lower shRNA electroporation efficiency. This possibility could be tested by inserting shRNA or siRNA into fertilized eggs by microinjection, instead of electroporation, which would be followed by assessing gene expression and phenotypes.

During *Clytia* early embryogenesis, *Wnt3* RNA is localized to the animal cortex of the egg and contributes to the formation of the oral-aboral axis^10^. Accordingly, knockdown of *Wnt3* in *Clytia* fertilized eggs induces suppression of axis formation and planula elongation (Fig. 4). In contrast, despite the fact that no morphological phenotype was observed after *Wnt3* knockdown in *Cladonema*, gene expression of the conserved axial patterning genes, *Brachyury* and *FoxQ2a*, was disrupted (Figs. 5G-5H and Supplementary Fig. 7). In addition to *Clytia*, the Wnt/β-catenin pathway is involved in axis formation during cnidarian embryogenesis including *Nematostella*^44,56,59^, *Acropora*^60^, *and Hydractinia*^57^. Furthermore, given that the Wnt/β-catenin pathway is also associated with axis formation during metamorphosis in *Hydractinia*^57,61^ and *Clytia*^62^, the reduced metamorphosis rate observed in *CpWnt3* knockdown *Cladonema* may be attributed to disrupted axis formation (Supplementary Table 5). Taken together, our data support the pivotal role of Wnt/β-catenin signaling in cnidarian development.

An interesting feature of jellyfish is that their morphology dramatically changes across their life cycle from planula to polyp and then to medusa. In particular, polyps and medusae, different adult stages of the same animal, exhibit distinct regenerative ability, lifespan, and behaviors. Although regeneration mechanisms have been extensively studied in polyp-only animals such as *Hydra* and *Hydractinia*, classical studies have used the medusa stage of several jellyfish species and demonstrated their regenerative potential^21^. Recent work using *Clytia* medusae has further shown a remarkable remodeling and repatterning mechanism orchestrated by muscle systems upon organ loss^17^. At the behavior level, the medusae of the upside-down jellyfish *Cassiopea* exhibit a sleep-like state^63^, which is similarly observed in *Hydra* polyps^64^. More recent work using transgenic *Clytia* has characterized feeding behaviors in medusae at a neural-network resolution^19^. To understand how these diverse biological processes in jellyfish are regulated at the molecular level, the functions of specific genes at various stages must be analyzed. Although gene knockdown by dsRNA electroporation into polyps in the scyphozoan jellyfish *Aurelia* has been reported^65^, gene knockdown at the medusa stage has not yet been achieved. Given that siRNA electroporation is applicable to *Hydra* polyps^33,66^ in addition to cnidarian fertilized eggs, we now have a better chance to apply this technique to the medusa stage, although, because they are susceptible to electric shock, electroporation parameters must be optimized in order to achieve gene knockdown in medusae. Collectively, our gene knockdown via siRNA electroporation method will complement the existing shRNA electroporation approach in cnidarian polyps and will enable molecular-level analysis of the vast biological phenomena exhibited across the different life stages in jellyfish.

## Acknowledgement

We thank R. Deguchi (Miyagi Univ. Education, Japan) for sharing *Cladonema pacificum* and EMBRC France for sharing *Clytia hemisphaerica*. We thank T. Momose (Sorbonne University, CNRS, France) for helpful discussion. We thank I. Nagai, H. Nakatani, and A. Sasaki for technical assistance; and A. Dahal, A. Tanimoto, and J. Higuchi for *Nematostella* culture. This work was supported by JST grant number JPMJCR1852 to E.K., AMED under grant number JP21gm6110025 to Y.N., and the JSPS KAKENHI grant numbers JP21H05255 to E.K., and JP17H06332, JP19K22550, JP22H02762 to Y.N.

## Author contributions

T.M.-O. conceptualized and designed the project, performed experiments using jellyfish, analyzed data, prepared figures, and wrote the manuscript. S.F. performed experiments and analyzed data. R.N. performed experiments using *Nematostella*. H.W. conceptualized and designed the project. E.K. contributed reagents. Y.N. conceptualized and designed the project, prepared figures, and wrote the manuscript. All authors approved the final manuscript.

## Data availability

The datasets used and/or analysed during the current study are available from the corresponding author on reasonable request and nucleotide sequences are available in the GenBank/EMBL/DDBJ under the accession numbers (*CpWnt1*, LC720432; *CpWnt1b*, LC720433; *CpWnt2*, LC720434; *CpWnt3*, LC720435; *CpWnt5*, LC720436; *CpWnt6*, LC720437; *CpWnt8*, LC720438; *CpWntA*, LC720439; *CpBrachyury*, LC720440; *CpFoxQ2a*, LC720441; *F-actin capping protein subunit beta*, LC720442; *CpEF1alpha*, LC720443).

