## Supplementary information for "siRNA-mediated gene knockdown via electroporation in hydrozoan jellyfish embryos"

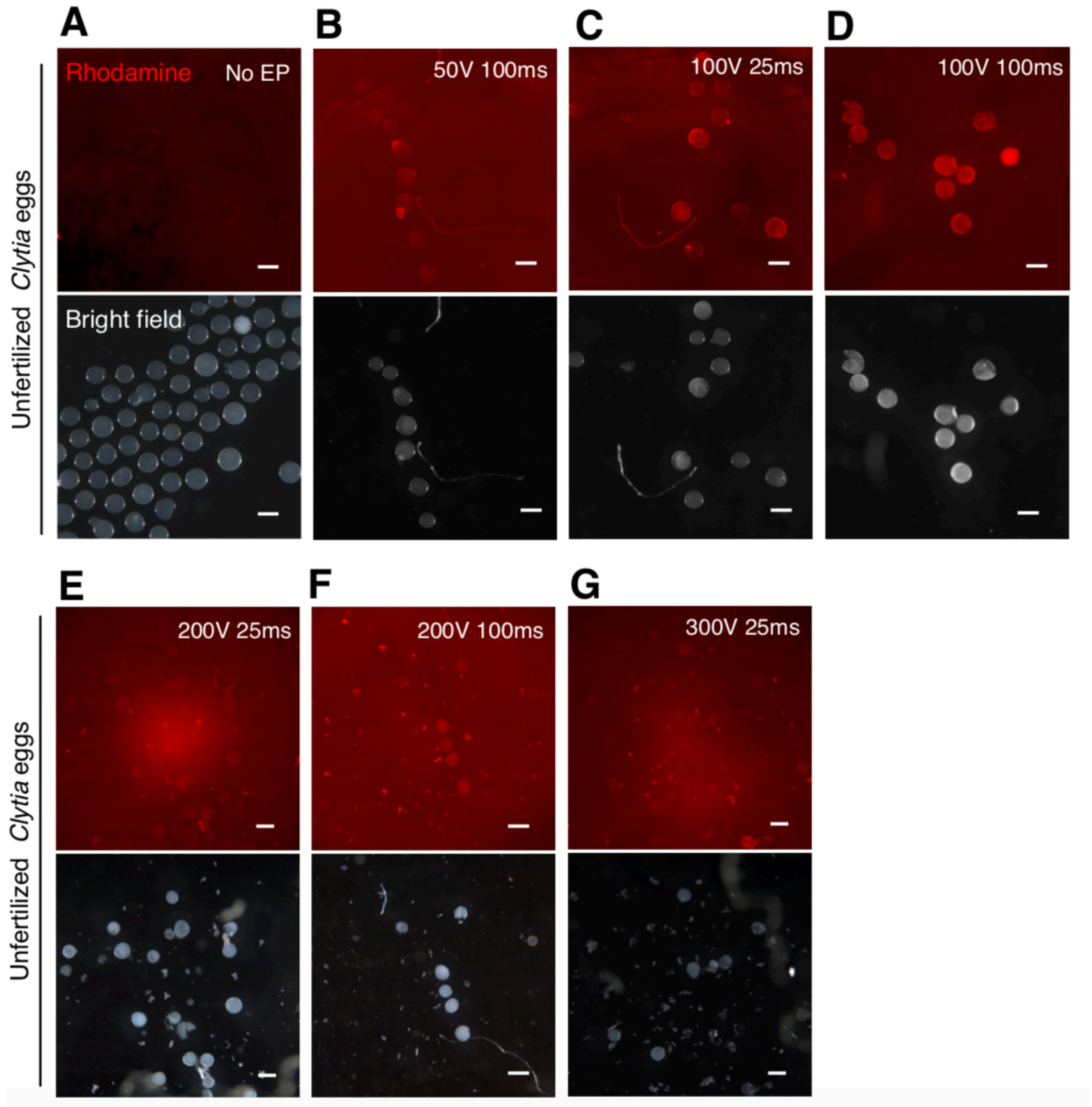

**Supplementary Figure 1. Dextran delivery to *Clytia* eggs under different electroporation conditions.** (**A**-**G**) Photos of unfertilized *Clytia* eggs after electroporation of Dextran-Rhodamine (1 mg/ml) in indicated conditions. Fluorescence intensity tends to increase in a voltage and time dependent manner. In the extreme conditions above 200 V (**E**-**G**), eggs were severely damaged. Scale bars: 200 μm.

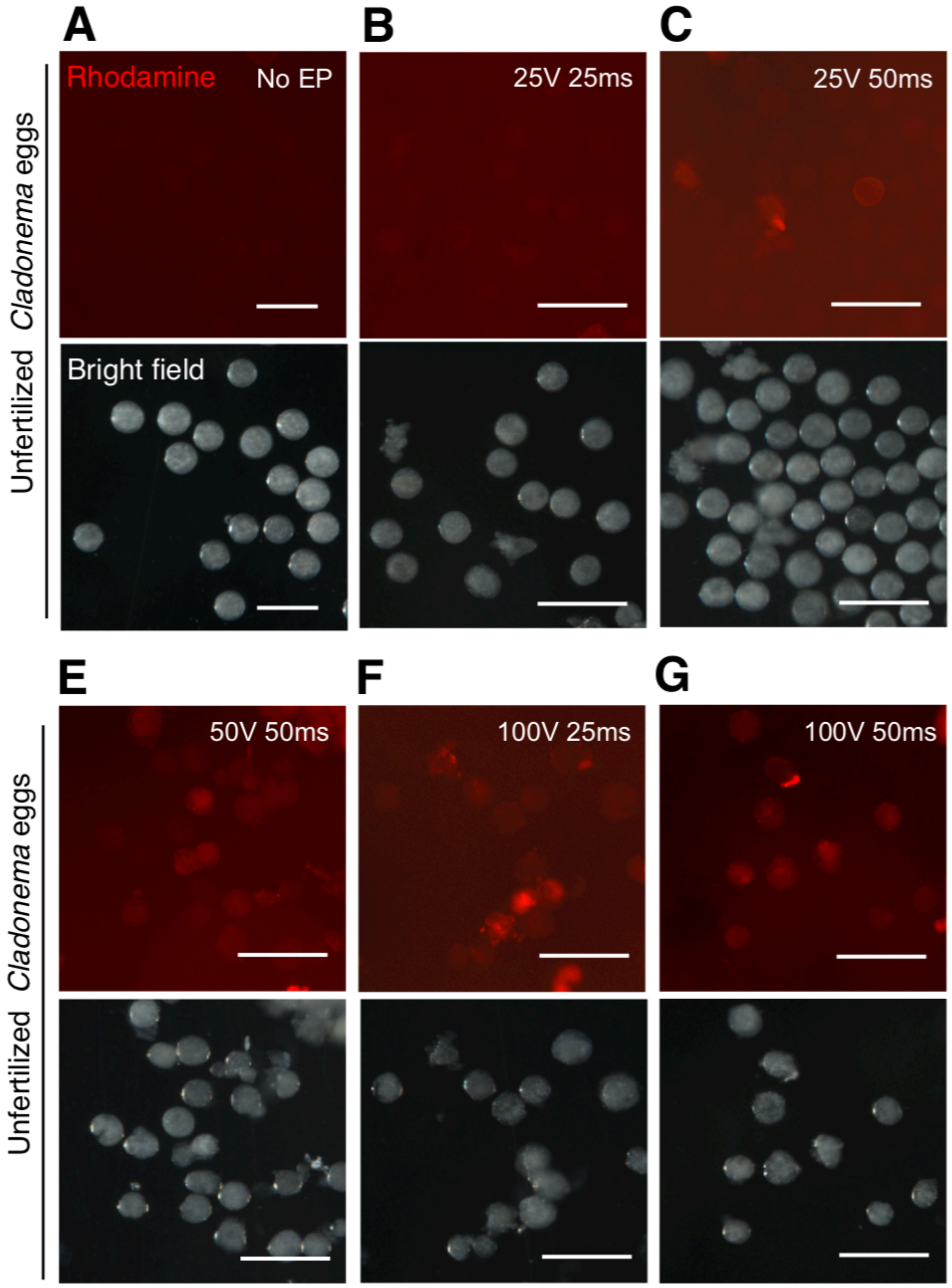

**Supplementary Figure 2. Dextran delivery to *Cladonema* eggs under different electroporation conditions.** (**A**-**F**) Photos of unfertilized *Cladonema* eggs after electroporation of Dextran-Rhodamine (1 mg/ml) in indicated conditions. Fluorescence was frequently detected beyond 50 V, but in the extreme conditions (**E** and **F**), eggs were severely damaged. Scale bars: 200 µm.

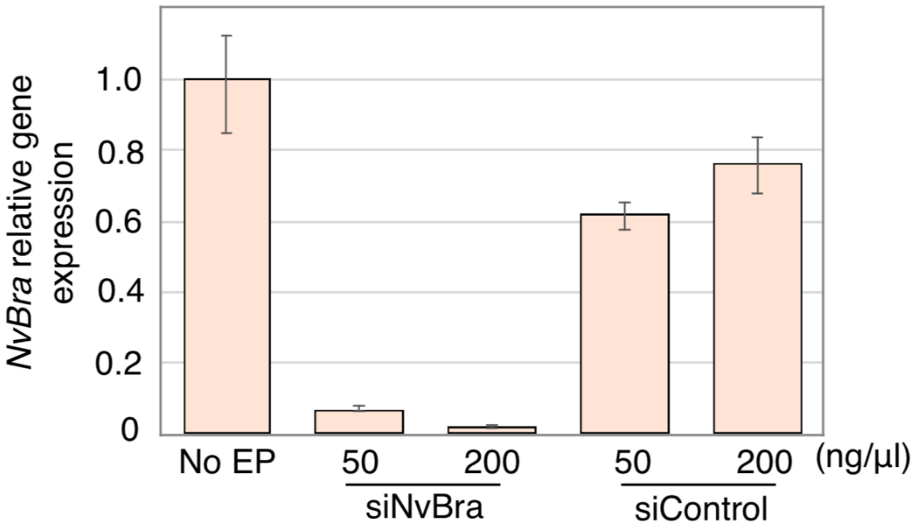

**Supplementary Figure 3. Gene knockdown with siRNA via electroporation of *Nematostella* fertilized eggs.** qPCR quantification of *Nematostella Brachyury* (*NvBra*) mRNA expression in siRNA-electroporated planulae (3 dpf). *NvEf1alpha* was used as an internal control. Shown are the results of the mean ± S.D. from a single experiment in triplicate, representing at least two independent experiments with similar results.

**A**

| Gene ID | CDS ID | memo | siCheGFP1_1 | siCheGFP1_2 | siCheGFP1_3 | CheGFP1 qP FW | CheGFP1 qP FW | CheGFP1_2 qP FW | CheGFP1_2 qP RV |
|---|---|---|---|---|---|---|---|---|---|
| | HQ397706.1 | CDS for siRNA design | C | C | C | C | C | C | C |
| XLOC_006257 | TCONS_00011161 | Expressed in planula | C | C | C | M1 | C | C | C |
| XLOC_043985 | TCONS_00070706 | Expressed in planula | C | M1 | M2 | C | | C | C |
| XLOC_010544 | TCONS_00018823 | Expressed in planula | M1 | M4 | M1 | | C | C | C |
| XLOC_035583a | TCONS_00056485 | | M4 | M5 | M2 | M1 | | C | M1 |
| XLOC_035583b | TCONS_00056484 | | M4 | M5 | M2 | M1 | | C | M1 |
| XLOC_035583c | TCONS_00056483 | | M4 | M5 | M2 | M1 | | C | M1 |
| XLOC_035557 | TCONS_00056425 | | C | C | M1 | M1 | M2 | M2 | C |

C, complete match. M, number of mismatch.

**B**

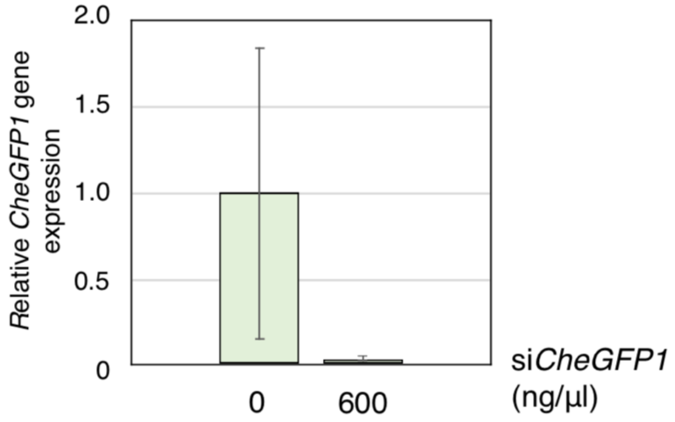

**Supplementary Figure 4. RT-qPCR analysis for the multiple *CheGFP1* loci in *Clytia*.** (**A**) A comparison of the seven *CheGFP1* loci and the previously published *CheGFP1* (HQ397706.1). The siRNA and PCR primers listed here are highlighted if they are expected to complement the *CheGFP1* gene (as C and M1). (**B**) Relative gene expression of *CheGFP1s*. RT-qPCR was conducted using the newly designed primer set that amplifies the seven *CheGFP1* genes. Error bars show standard error. Experiments were performed in triplicate and repeated three times. p= 0.376.

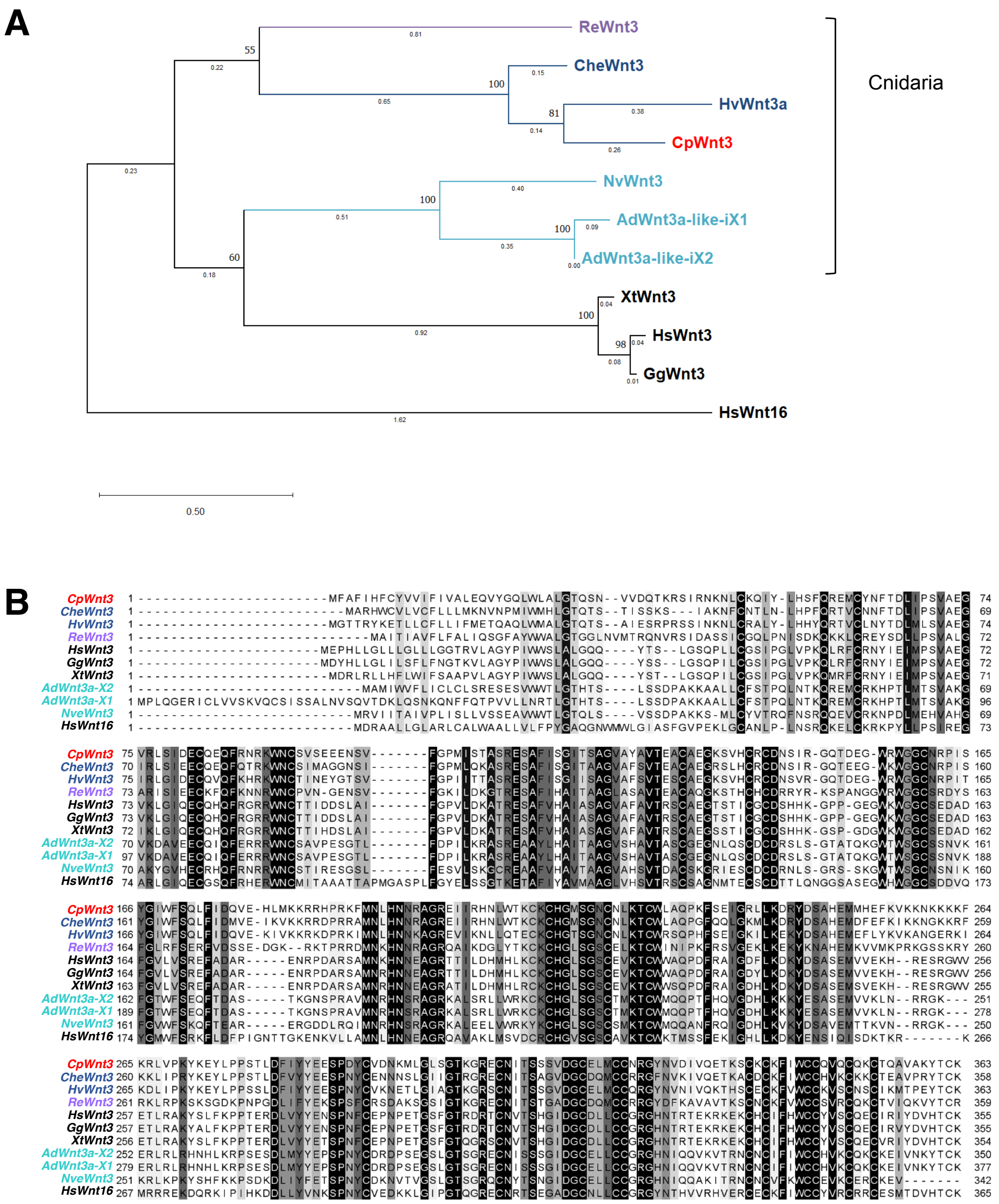

**Supplementary Figure 5. Identification of *Cladonema* Wnt3 by phylogenetic analyses.** The results of (A) phylogenetic analyses and (B) sequence alignment of Wnt3 among indicated species. HsWnt16 was used as an outgroup. *Cladonema* Wnt3 (CpWnt3) is highlighted in red. Other cnidarian Wnt3s are highlighted in other colors (Schyphozoa: purple, Hydrozoa: midnight blue, Anthozoa: turquoise). (B) In the sequence alignment, consensus residues are indicated with black and grey. CpWnt3 protein showed conserved amino acid sequence with Wnt3 proteins of other species. *Re*: *Rhopilema esculentum*, *Che*: *Clytia hemisphaerica*, *Hv*: *Hydra vulgaris*, *Cp*: *Cladonema pacificum*, *Nv*: *Nematostella vectensis*, *Ad*: *Acropora digitifera*, *Xt*: *Xenopus tropicalis*, *Hs*: *Homo sapiens*, *Gg*: *Gallus gallus*. GenBank accession numbers are listed in Supplementary Table 3.

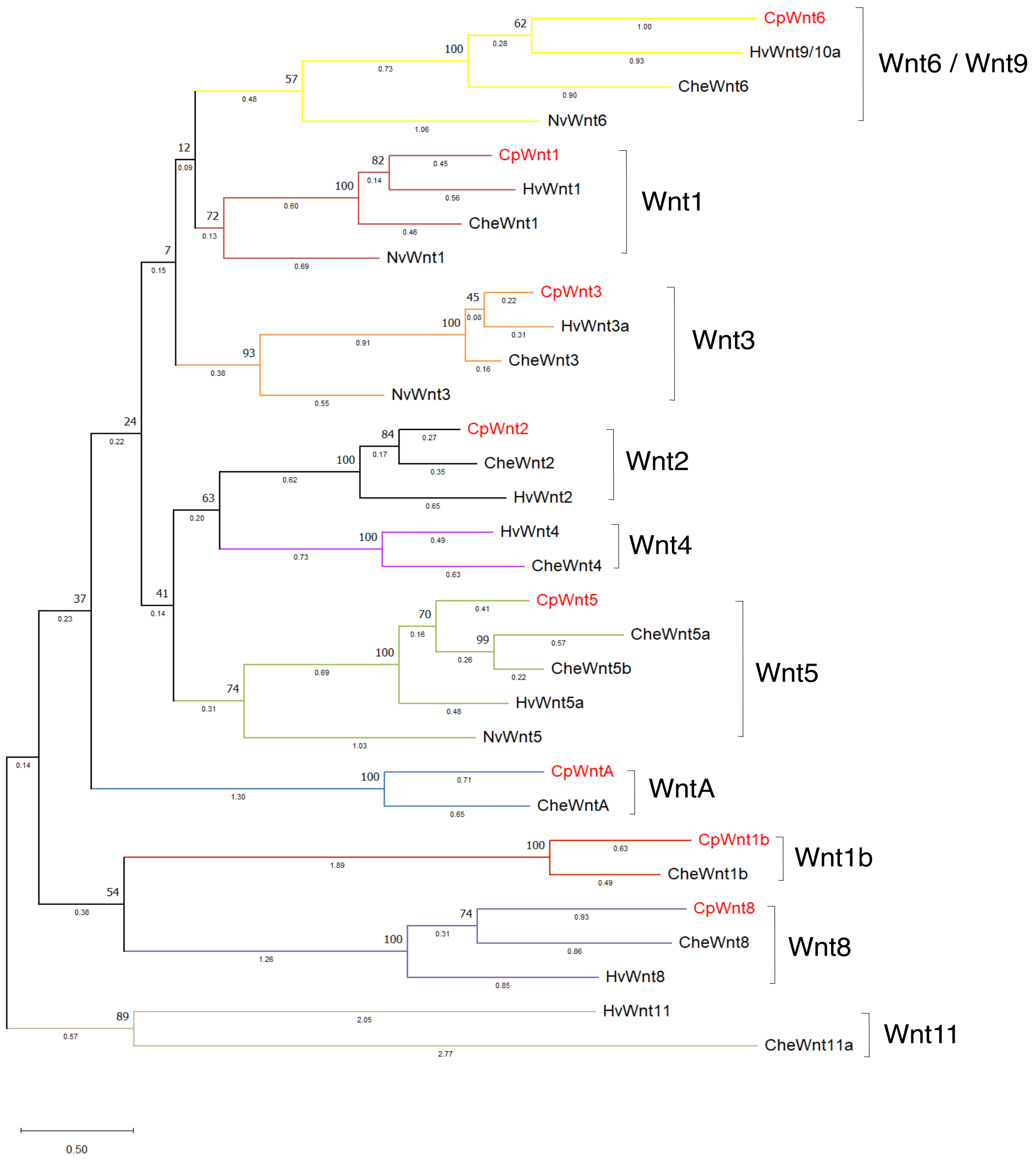

**Supplementary Figure 6. Phylogenetic analysis of *Cladonema* Wnt proteins.**

To analyze the orthology of *Cladonema* Wnt proteins, phylogenetic analysis was performed using the amino acid sequences of Wnt proteins from *Cladonema* (Cp), *Clytia* (Che), *Hydra* (Hv), and *Nematostella* (Nv). We performed *Cladonema* Wnt ligand CDS annotation using the *Clytia* CDS database (MARIMBA, Marine models database: http://marimba.obs-vlfr.fr) and found eight contigs annotated with Wnt ligands in *Cladonema*. GenBank accession numbers and transcriptome name or RNAseq contig ID are listed in Supplementary Table 4.

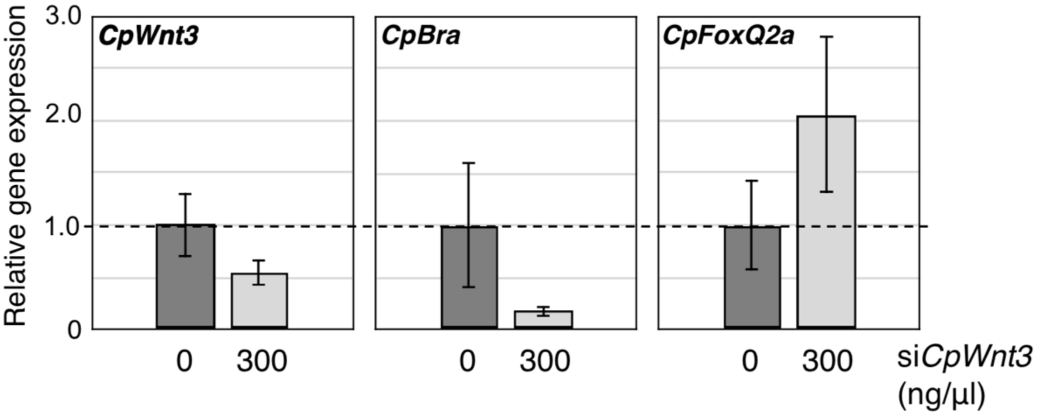

**Supplementary Figure 7. qPCR quantification of axial marker genes *in Cladonema* planulae.** Quantification of *CpWnt3*, *CpBra*, and *CpFoxQ2a* mRNA levels of 1-day planulae (0 ng or 300 ng/µl of si*CpWnt3*) by RT-qPCR. The *beta-actin* gene was used as an internal control. Bar heights represent mean values. Gene expression levels are standardized relative to control (0 ng/µl). Error bars show standard error. n=8. p-values: *CpWnt3*, p=0.213; *CpBra*, p=0.224; *CpFoxQ2a*, p=0.224.

| | 1 hpf survival (%) | SD | *n* |
|---|---|---|---|
| No EP | 59.67 | 17.01129 | 338 |
| Fertilized egg EP | 60.15 | 14.64567 | 403 |
| Unfertilized egg EP | 7.04 | 0.907342 | 672 |

* Average of at least 3 biologically different experiments

**Supplementary Table 1. Survival rate of *Cladonema* embryos after electroporation.**
The number of cleavage stage *Cladonema* embryos at 1 h post fertilization (hpf) was considered embryo survival. Compared to fertilized eggs, unfertilized eggs were much less tolerant of electroporation.

| | siRNA ng/µL | Survival rate 3 dpf (%) | n |
|---|---|---|---|
| No electroporation (just dejellied) | 0 | 40 | 610 |
| *NvBrachyury* siRNA | 50 | 24 | 696 |
| *NvBrachyury* siRNA | 200 | 32 | 383 |
| Negative control siRNA | 50 | 32 | 398 |
| Negative control siRNA | 200 | 32 | 491 |

**Supplementary Table 2**. **Survival rate of *Nematostella* embryos after electroporation.** The number of live *Nematostella* larvae was counted at 3 dpf (planula stage). The survival rate remained around 20-40% regardless of siRNA concentration or target.

| Protein name | Protein or Nucleotide ID | |
|---|---|---|
| **HsWnt3** | BAB70502.1 | *Homo sapiens* |
| **GgWnt3** | ABK90822.1 | *Gallus gallus* |
| **XtWnt3** | ABG49501.1 | *Xenopus tropicalis* |
| **CpWnt3** | LC720435 | *Cladonema pacificum* |
| **CheWnt3** | ACB15465.1 | *Clytia hemisphaerica* |
| **NvWnt3** | ABF48092.1 | *Nematostella vectensis* |
| **AdWnt3a-like isoform X2** | XP_015753662.1 | *Acropora digitifera* |
| **AdWnt3a-like isoform X1** | XP_015753659.1 | *Acropora digitifera* |
| **HvWnt3a** | QCF59212.1 | *Hydra vulgaris* |
| **ReWnt3** | AID70699.1 | *Rhopilema esculentum* |
| **WNT16** | AAD49351.1 | *Homo sapiens* |

**Supplementary Table 3. Wnt3 proteins in Supplementary Figure 5.**

GenBank accession numbers and transcriptome name or RNA-seq contig ID of *Wnt3*s and *HsWNT16* are shown.

| Protein name | gene ID | protein ID | Old name |
|---|---|---|---|
| **CheWnt1** | XLOC_015404 | TCONS_00027371 | CheWntX2 |
| **CheWnt1b** | XLOC_015382 | TCONS_00027333 | - |
| **CheWnt2** | XLOC_031686 | TCONS_00050019 | CheWntX1A |
| **CheWnt3** | XLOC_001931 | TCONS_00003473 | CheWnt3 |
| **CheWnt4** | XLOC_041060 | TCONS_00065525 | CheWntX1B |
| **CheWnt5** | XLOC_000650 | TCONS_00001198 | CheWnt5 |
| **CheWnt5b** | XLOC_030003 | TCONS_00047033 | - |
| **CheWnt6** | XLOC_035748 | TCONS_00056741 | CheWnt9 |
| **CheWnt8** | XLOC_045775 | TCONS_00073720 | - |
| **CheWnt11a** | XLOC_002938 | TCONS_00005153 | CheWntX3 |
| **CheWntA** | XLOC_006909 | TCONS_00012270 | - |
| | | **Nucleotide ID** | |
| **CpWnt1** | | LC720432 | |
| **CpWnt1b** | | LC720433 | |
| **CpWnt2** | | LC720434 | |
| **CpWnt3** | | LC720435 | |
| **CpWnt5** | | LC720436 | |
| **CpWnt6** | | LC720437 | |
| **CpWnt8** | | LC720438 | |
| **CpWntA** | | LC720439 | |
| | | **Protein ID** | |
| **HvWnt1** | | BAH23782.1 | |
| **HvWnt2** | | BAH23783.1 | |
| **HvWnt3** | | QCF59212.1 | |
| **HvWnt4** | | XP_047138590.1 | |
| **HvuWnt5** | | BAH23774.1 | |
| **HvWnt8** | | BAH23780.1 | |
| **HvWnt9/10a** | | BAH23777.1 | |
| **HvWnt11** | | BAH23776.1 | |
| | | **Protein ID** | |
| **NvWnt1** | | AAT00640.1 | |
| **NvWnt3** | | ABF48092.1 | |
| **NvWnt5** | | AAW28133.1 | |
| **NvWnt6** | | AAW28134.1 | |

**Supplementary Table 4**. **Wnt family genes in Supplementary Figure 6.**

This table shows GenBank accession numbers and transcriptome name or RNA-seq contig ID of Wnt ligands. Protein names of Wnt ligands of *Clytia* correspond to the new names given by Condamine et al[15].

| | Egg to planula | | Planula to polyp | | |
|---|---|---|---|---|---|
| | 1 dpf planula† | SE | Metamorphosis rate (%) ‡ | SE | n§ |
| Control (0 ng/μl) | 206.33 | 91.26 | 20.68 | 3.39 | 472 |
| si*CpWnt3* (300 ng/μl) | 139.50 (n.s.) | 94.92 | 9.36 (*) | 2.07 | 434 |

† Average number of planulae (1 dpf) from 6 different experiments
‡ Average rate of metamorphosis (7dpf) of 3 biologically different experiments
§ Total number of planulae (1 dpf)
n.s., not significant, *p<0.05.

**Supplementary Table 5**. **Survival rate of *Cladonema* embryos and metamorphosis rate**. The number of planulae (1 dpf) and percentage of metamorphosis from planula to polyp. *CpWnt3* knockdown planulae showed significant decline in metamorphosis rate. p=0.046.

### Protocol: siRNA electroporation for hydrozoan jellyfish embryos

**Materials**

- Ficoll (M.W.400,000 EP (Extra Pure Reagent); Nacalai tesque; Cat No.16006-92)
- Artificial sea water (ASW; *Clytia*; 220g SEA LIFE per 5 L MilliQ water, *Cladonema*; 24 p.p.t)
- RNase free water (Nippon gene; supplied with the siRNA)
- V7 cup (φ63 × φ58 × 34.6 mm; AS ONE; Cat No. 5-067-27)
- 60 mm dish (60 x 15 mm; Corning; Cat No. 430166)
- Electroporation cuvette (4 mm; Bio-Rad; Cat No. 1652088)
- Gene Pulser Xcell complete system (Bio-Rad; Cat No. 1652660J1)

Note:

- 15% Ficoll in ASW was prepared according to previous reports (Karabulut et al., 2019 *Dev Biol* and Quiroga-Artigas et al., 2020 *Sci Rep*).
- 1.5ml microtubes, 200 µl pipet tips and 6 mm dishes were coated with 15% Ficoll prior to preparing electroporation samples.

#### Embryo preparation and electroporation protocol

The following processes are conducted at room temperature (20-22°C).

1-a. *Clytia*: After light stimulation (in 45-60 min), transfer medusae into a V7 cup.

1-b. *Cladonema*: Before dark stimulation, transfer male medusae into 60 mm dishes (5-10 medusae per one dish). Transfer female medusae into a V7 cup.

2. After spawning, collect unfertilized eggs into 60 mm dish (*Clytia*) or 1.5 ml microtubes (*Cladonema*) using Ficoll-coated 200 µl pipet tips (cut off the end of the tip to make a larger hole).

3-a. *Clytia*: Add sperm water into unfertilized eggs in dish and incubate for 5 min for fertilization. Carefully transfer eggs into Ficoll-coated 1.5 ml microtube.

3-b. *Cladonema*: After unfertilized eggs are settled, add sperm water into the microtube and then incubate for 5 min for fertilization.

4. Remove ASW from the fertilized egg solution.

5. Add 15% Ficoll in ASW to adjust the total volume for the number of electroporations (90 µl per each electroporation), and carefully mix them by pipetting.

6. Divide fertilized eggs into clean the Ficoll-coated 1.5 ml microtubes (90 µl each).

7. Add 10 µl of siRNA or shRNA solution, which is adjusted to the proper concentration in RNase free, to fertilized eggs, and carefully mix by pipetting.

   Note: RNA stock solutions are stored at -20°C. After thawing, keep the RNA mixture on ice throughout the experiment.

8. Transfer the mixture of fertilized eggs and RNA (total volume 100 µl) into a Ficoll-coated 4 mm cuvette using 200 µl pipet tip.

9. Insert the cuvette into the Gene Pulser Xcell Shockpod (Bio-Rad). Choose program No.3 (Square pulse) and set an experimental condition as 50 V, 25 msec, 4 mm. Press the red button to add pulse.

10. After 1 min-waiting, gently transfer the entire reaction solution into a 6 mm dish. Incubate it on the dish for 10 min.

11. Carefully remove 15% Ficoll in ASW and RNA solution and then add new ASW. Leave the embryos undisturbed for several hours until they recover.
